# Testing for differences in polygenic scores in the presence of confounding

**DOI:** 10.1101/2023.03.12.532301

**Authors:** Jennifer Blanc, Jeremy J. Berg

## Abstract

Polygenic scores have become an important tool in human genetics, enabling the prediction of individuals’ phenotypes from their genotypes. Understanding how the pattern of differences in polygenic score predictions across individuals intersects with variation in ancestry can provide insights into the evolutionary forces acting on the trait in question, and is important for understanding health disparities. However, because most polygenic scores are computed using effect estimates from population samples, they are susceptible to confounding by both genetic and environmental effects that are correlated with ancestry. The extent to which this confounding drives patterns in the distribution of polygenic scores depends on patterns of population structure in both the original estimation panel and in the prediction/test panel. Here, we use theory from population and statistical genetics, together with simulations, to study the procedure of testing for an association between polygenic scores and axes of ancestry variation in the presence of confounding. We use a general model of genetic relatedness to describe how confounding in the estimation panel biases the distribution of polygenic scores in a way that depends on the degree of overlap in population structure between panels. We then show how this confounding can bias tests for associations between polygenic scores and important axes of ancestry variation in the test panel. Specifically, for any given test, there exists a single axis of population structure in the GWAS panel that needs to be controlled for in order to protect the test. Based on this result, we propose a new approach for directly estimating this axis of population structure in the GWAS panel. We then use simulations to compare the performance of this approach to the standard approach in which the principal components of the GWAS panel genotypes are used to control for stratification.

**Author Summary:** Complex traits are influenced by both genetics and the environment. Human geneticists increasingly use polygenic scores, calculated as the weighted sum of trait-associated alleles, to predict genetic effects on a phenotype. Differences in polygenic scores across groups would therefore seem to indicate differences in the genetic basis of the trait, which are of interest to researchers across disciplines. However, because polygenic scores are usually computed using effect sizes estimated using population samples, they are susceptible to confounding due to both the genetic background and the environment. Here, we use theory from population and statistical genetics, together with simulations, to study how environmental and background genetic effects can confound tests for association between polygenic scores and axes of ancestry variation. We then develop a simple method to protect these tests from confounding, which we evaluate, alongside standard methods, across a range of possible situations. Our work helps clarify how bias in the distribution of polygenic scores is produced and provides insight to researchers wishing to protect their analyses from confounding.

## 1 Introduction

The calculation of polygenic scores [1] has become a routine procedure in many areas of human genetics. The promise of polygenic scores is that they provide a means for phenotypic prediction from genotype data alone. By measuring the association between a genetic variant and phenotype in a genome wide association study (GWAS), we get an estimate of its effect on the phenotype, averaged over the environments experienced by the individuals in that sample. These effect estimates can then be combined into polygenic scores in a separate prediction panel by taking a sum of the genotypes of individuals in that panel, weighted by the estimated effects. Under the relatively strict assumptions that genetic and environmental effects combine additively, that variation in the phenotype is not correlated with variation in ancestry within the GWAS panel, and that the prediction panel individuals experience a similar distribution of environments to the GWAS panel individuals, these scores can be viewed as an estimate of each individual’s expected phenotype, given their genotypes at the included sites. If these assumptions are met, polygenic scores would seem to provide a means of separating out at least some of the genetic effects on a given phenotype.

However, this promise of polygenic scores is also one of their main pitfalls. The effects of individual variants are typically estimated from population samples in which the environments that individuals experience vary as a function of their social, cultural, economic, and political contexts. Differences in these factors are often correlated with differences in ancestry within population samples, and these ancestry-environment correlations can induce systematic biases in the estimated effects of individual variants. Similar biases can also arise if genetic effects on the phenotype vary as a function of ancestry within the GWAS sample. Ancestry stratification is a long recognized problem in the GWAS study design [2], and many steps have been taken to guard against its effects. These include bias avoidance approaches, like the sampling of GWAS panels that are relatively homogeneous with respect to ancestry, and statistical bias correction approaches, such as the inclusion of genetic principal components as covariates [3], linear mixed models [4, 5], and LD score regression [6]. These approaches have largely been successful in minimizing the number of false positive single variant associations [7]. However, effect size estimates can still exhibit slight stratification biases that are not large enough to significantly alter the false discovery rates for individual variants, and these biases can be compounded when aggregating across loci, leading to confounded predictions in which the ancestry associated effects are mistaken for genetic effects.

Separation of direct genetic effects from correlations between ancestry and either the environment or the genetic background is important to all applications of polygenic scores. Empirically, polygenic scores exhibit geographic clustering even in relatively homogeneous samples and after strict control for population stratification [8, 9, 10, 11]. It is natural to ask if these observed differences reflect a real difference in the average genetic effect on the trait. From a population biology perspective, these patterns may be signals of natural selection [12] or phenotype biased migration [9]. Medically, it is interesting to know if polygenic score differences or gradients represent real underlying gradients in the average genetic effect [13], whether those gradients are caused by non-neutral evolutionary mechanisms or not. However, observed patterns of polygenic scores may also be driven by residual bias in effect size estimates, and stratification biases remain a persistent issue.

This issue has been particularly apparent in the detection of directional selection acting on complex traits. Polygenic scores are an ideal tool for this task, as studying the distribution of scores among individuals who differ in ancestry allows us to aggregate the small changes in allele frequency induced by selection on a polygenic trait into a detectable signal [14, 15, 16, 17]. Several research groups have developed and applied methods to detect these signals [18, 12, 19, 20, 21, 22, 23, 24]. However, these efforts have been met with challenges, as several papers reported signals of recent directional selection on height in Europe using effects obtained from GWAS meta-analyses [25, 26, 18, 12, 27, 28, 29, 20, 30, 31, 19], only for these signals to weaken substantially or disappear entirely when re-evaluated using effects estimated in the larger and more genetically homogeneous UK Biobank [32, 33, 22, 23]. Further analysis suggested that much of the original signal could be attributed to spurious correlations between effect size estimates and patterns of frequency variation, presumably induced by uncorrected ancestry stratification in the original GWAS [32, 33].

Recently, in the context of selection tests, Chen et al. [34] proposed a strategy to mitigate the impact of stratification by carefully choosing the GWAS panel so that even if residual stratification biases in effect size estimates exist, they will be unlikely to confound the test (see also [35] for examples of this approach). They reasoned that because polygenic selection tests ask whether polygenic scores are associated with a particular axis of population structure in a given test panel, and because the bias induced by stratification in effect sizes depends on patterns of population structure in the GWAS panel [27], then one should be able to guard against bias in polygenic selection tests by choosing GWAS and test panels where the patterns of population structure within the two panels are not expected to overlap.

However, this approach comes at a cost of reduced power: polygenic scores are generally less accurate when the effect sizes used to compute them are ported to genetically divergent samples [36, 37, 38, 39, 40]. Less accurate polygenic scores are then less able to capture evolution of the mean polygenic score, all else equal [39]. These decays in polygenic score accuracy also pose a significant challenge to their use in medicine, as scores that are predictive for some and not for others may exacerbate health inequities [41]. Thus, realizing the potential of polygenic scores in both basic science and medical applications will require the use of large and genetically diverse GWAS panels. Successfully deploying polygenic scores developed from these diverse panels will require that we have a precise understanding of how bias is produced in polygenic score predictions, and the development and evaluation of methods to protect against this bias.

In this paper, we first model the covariance of genotypes in a GWAS and test panel in terms of an underlying population genetic model, and give expressions for the bias in the distribution of polygenic scores as a function of the underlying model. We then show how bias in the association between polygenic scores and a specific axis of ancestry variation in the test panel depends on the extent to which potential confounders in the GWAS lie along a specific axis of ancestry variation in the GWAS panel. Next, we evaluate ways to control for confounding along this axis, including the standard PCA-based approach, as well as a new approach that uses test panel genotypes to estimate the axis directly. We find that the utility of each approach depends on a host of factors, including the number of independent SNPs used to compute the correction, the number of samples in the GWAS panel, and the amount of variance in the GWAS panel explained by the target axis.

## 2 Model

To model the distribution of genotypes in both panels, we assume that each individual’s expected genotype at each site can be modeled as a linear combination of contributions from a potentially large number of ancestral populations, which are themselves related via an arbitrary demographic model. Natural selection, genetic drift, and random sampling each independently contribute to the distribution of genotypes across panels, and we make the approximation that these three effects can be combined linearly. In supplemental section S1 we develop the full population model which we then extend to individuals. In the main text, we present just the individual genotype model, along with our model of the phenotype.

### 2.1 Genotypes

We consider two samples of individuals, one to compose the GWAS panel and one to compose the test panel. Individuals in each panel are created as mixtures of an arbitrary number of *K* underlying populations related via an arbitrary demographic model (see supplement section S1.1 and S1.2), where *a*_*𝓁*_ is the ancestral allele frequency at site 𝓁. There are *N* test panel individuals and the vector of deviations of their genotypes from the mean genotype in the ancestral population (2*a*_*𝓁*_) is

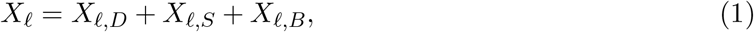

where *X*_*𝓁,D*_ and *X*_*𝓁,S*_ are the deviations due to drift and natural selection, respectively. We can think of the quantity 2*a*_*𝓁*_ + *X*_*𝓁,D*_ + *X*_*𝓁,S*_ as giving a set of expected genotypes given the evolutionary history of the populations from which the test panel individuals were sampled from, while *X*_*𝓁,B*_ contains the binomial sampling deviations across individuals given these expected genotypes.

Similarly, for the *M* GWAS panels individuals, the deviation of their genotypes can be decomposed as

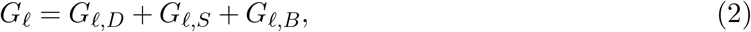

where *G*_*𝓁,D*_ and *G*_*𝓁,S*_ are the deviations due to drift and selection. *G*_*𝓁,B*_ captures the binomial sampling variance given the expected genotypes of the GWAS panel individuals.

Individuals in the two panels may draw ancestry from the same populations, or from related populations, which induces the joint covariance structure

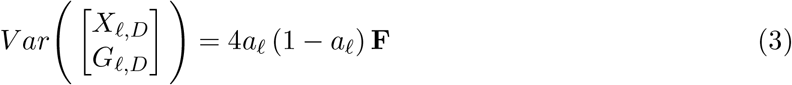

where the matrix

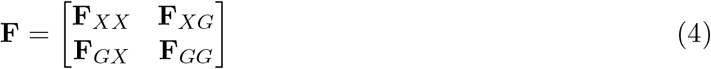

contains the within and between panel relatedness coefficients. Entries of **F** give the relatedness between pairs of individuals given the underlying demographic model and the fraction of ancestry each individual draws from each population. As such, the entries of **F** are directly related to the expected pairwise coalescent times between pairs of samples, given the demographic model [42].

### 2.2 Phenotypes

We assume that individuals in the GWAS panel are phenotyped and that the trait includes a contribution from *S* causal variants, which make additive genetic contributions, as well as an independent environmental effect. The vector of mean-centered phenotypes for the *M* individuals in the GWAS panel can then be written

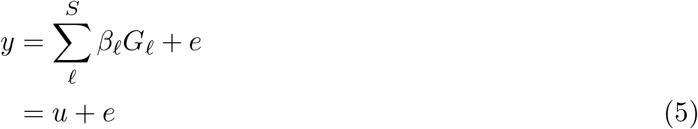

where 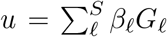 is the combined genetic effect of all *S* causal variants, and *e* represents the combination of all environmental effects.

We assume that the environmental effect on each individual is an independent Normally distributed random variable with variance 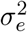, but that the expected environmental effect can differ in some arbitrary but unknown way across individuals. We write the distribution of environmental effects as 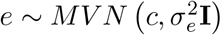, where *c* is the vector of expected environmental effects.

Similar to our decomposition in eq. 2, the genetic effect, *u*, can be broken down into the contributions from drift, selection, and binomial sampling such that *u* = *u*_*D*_ +*u*_*S*_ +*u*_*B*_. Here 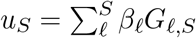 contains fixed effects reflecting the expected genetic contributions to the phenotype, given history of selection acting on the phenotype, and given the ancestries of the individuals in the GWAS panels (see supplement section S1.4). Both *u*_*D*_ and *u*_*B*_ have expectation zero, so 𝔼[*u*] = *u*_*S*_. The vector of individuals’ expected phenotypes, given their ancestry and socio-environmental contexts, is therefore given by *u*_*S*_ + *c*. We assume that these are not known.

## 3 Results

Now, given these modeling assumptions, we describe how the relationship between the GWAS and test panels impacts the distribution of polygenic scores and the association between the polygenic scores and a given axis of population structure which is observed only in the test panel. We first consider the case where no attempt is made to correct for population structure. Motivated by these results, we then outline the conditions that need to be met in order to ensure an unbiased association test. Finally, we explore how two different correction strategies, the standard PCA approach and a novel approach that uses the test panel genotypes, play out in practice.

### 3.1 The impact of stratification bias on polygenic scores

We consider a vector of mean centered polygenic scores, computed in the test panel. If the causal effects (*β*_*𝓁*_) were known, then the polygenic scores would be given by

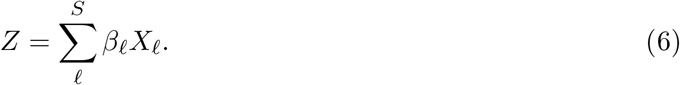

Of course, the causal effects are not known, and must be estimated in the GWAS panel. Conditional on the genetic and environmental effects on the phenotypes of the individuals in the GWAS panel (i.e. *u* and *e*), and genotypes at the focal site (*G*_*𝓁*_), the marginal effect size estimate for site 𝓁 is given by

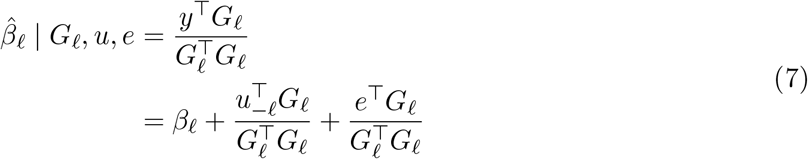

where we have decomposed the genetic effect into the causal contribution from the focal site and the contribution from the background, i.e. *u* = *β*_*𝓁*_*G*_*𝓁*_ + *u*_−*𝓁*_. This allows us to further decompose the marginal association in eq. 7 into the causal effect (*β*_*𝓁*_), the association between the focal site and the background genetic contribution from all other sites 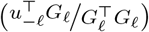, and the association with the environment 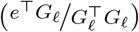.

The deviation of an allele’s estimated effect size from its expectation depends in part on *G*_*𝓁,D*_, the component of variation in the GWAS panel genotypes due to genetic drift. Because *G*_*𝓁,D*_ can be correlated with *X*_*𝓁,D*_ (deviations due to drift in test panel genotypes) due to shared ancestry, the estimated effect sizes can become correlated with the pattern of genotypic variation in the test panel for reasons that have nothing to do with the actual genetic effect of the variant. This leads to a bias in the polygenic scores,

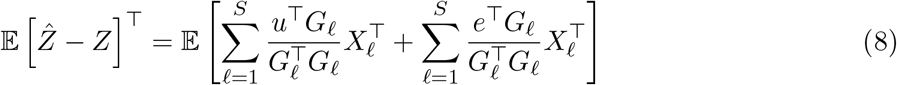

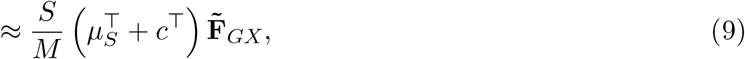

(see section S3) where *µ*_*S*_ is the vector of expected genetic backgrounds, *c* is the vector of expected environmental effects, and

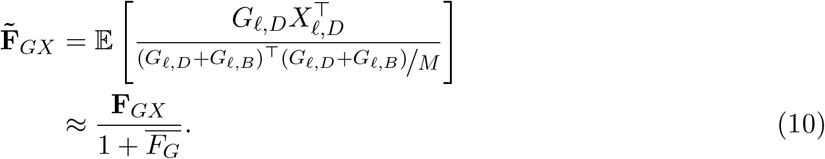

Here 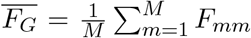 is the average level of self relatedness in the GWAS panel and 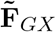 is the expected cross-panel genetic relatedness matrix computed on standardized genotypes, which is approximately equal to 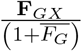 if 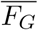 is small.

If the GWAS and test panels do not overlap in population structure, then 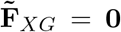, and the polygenic scores are unbiased with respect to ancestry (i.e.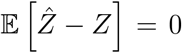), independent of the confounders, *µ*_*S*_ and *c* [1, 34, 35]. Stratification may still bias individual effects, but these residual biases are indistinguishable from noise from the perspective of the polygenic scores, as they are uncorrelated with all axes of population structure present in the test panel.

### 3.2 Bias in polygenic scores leads to biased polygenic score associations

We want to test the hypothesis that the polygenic scores are associated with some test vector, *T*. We assume that *T* is measured only in the test panel, and might represent an eco-geographic variable of interest (e.g latitude [12] or an encoding of whether one lives in a particular geographic region or not [9, 43], the fraction of an individual’s genome assigned to a particular “ancestry group”[18, 20], or one of the top genetic principal components of the test panel genotype matrix [21]).

To test for association of polygenic scores with the test vector, we take our test statistic the as slope of the regression of the polygenic scores against the test vector, which we denote *q*. Assuming *T* is standardized, this slope is given by

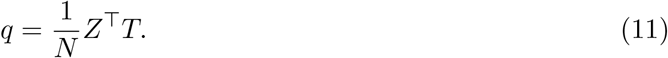

A more powerful test is available by modeling the neutral correlation structure among individuals due to relatedness (see section S8), but the simpler i.i.d. model presented here is sufficient for our purposes. Under the null model where selection has not perturbed allele frequencies in the test panel, 𝔼[*q*] = 0, reflecting the fact that genetic drift is directionless.

In practice, an estimate of *q* is obtained using the polygenic scores computed from estimated effect sizes, i.e. 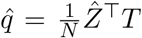. The bias in the polygenic score association test statistic 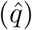 then follows straightforwardly from the bias in the polygenic scores,

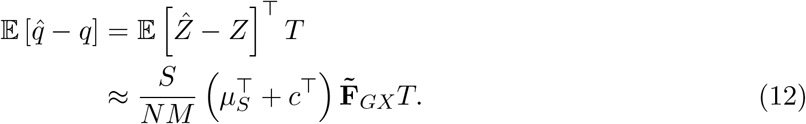

Therefore, we expect the polygenic score association test to be biased when the test vector (*T*) aligns with the vector of expected phenotypes (*µ*_*S*_ + *c*) in a space defined by the cross panel genetic similarity matrix 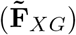. The conditions for an unbiased polygenic score association test are therefore narrower than the conditions needed to ensure unbiased polygenic scores in general. Rather than requiring that 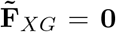, we need only to ensure that certain linear combination of the entries of 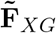 are equal to zero, i.e. that 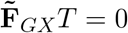.

We can gain further intuition by expressing the association statistic, *q*, in a different way. Specifically, we can re-frame this test as a statement about the association between the effect sizes and a set of genotype contrasts, 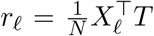, which measure the association between the test vector and the genotypes at each site [12]. Writing *β* and *r* for the vectors of effect sizes and genotype contrasts across loci, the association test statistic can be rewritten as

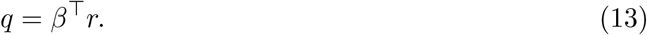

This allows us to rewrite the bias in the estimator, 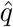, as

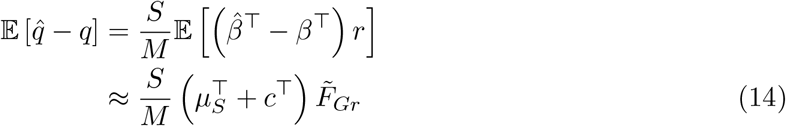

where

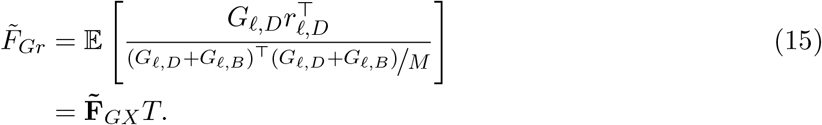

Here eq. 14 expresses the bias entirely in terms of vectors that belong to the GWAS panel: for each GWAS panel individual *m*, 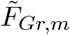 measures the covariance between individual *m*’s genotype and the genotype contrasts of the test, standardized at each site by the variance of genotypes across individuals in the GWAS panel (eq. 15). Thus, 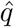 is biased when the vector of expected phenotypes (*µ*_*S*_ +*c*) aligns with this vector of standardized covariances 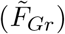. Confounders which are orthogonal to this axis do not generate bias in the association test, even if they bias the polygenic scores along other axes.

### 3.3 Controlling for stratification bias in polygenic association tests

Given the above results, how can we ensure that patterns we observe in the distribution of polygenic scores are not the result of stratification bias? As discussed above, a conservative solution is to prevent bias by choosing a GWAS panel that does not have any overlap in population structure with the test panel, but this is not ideal due to the well documented portability issues that plague polygenic scores [36, 44, 40], and because it limits which GWAS datasets can be used to test a given hypothesis. Another obvious solution is to include the vectors of expected genetic and environmental effects, *u*_*S*_ and *c* respectively, as covariates in the GWAS. Doing so would remove all ancestry associated bias from the estimated effects, and thus ensure that any polygenic score association test carried out using these effects would be unbiased. However, *u*_*S*_ and *c* are typically not measurable, so this is generally not an option. Alternatively, our analysis above suggests that including 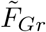 as a covariate in the GWAS model is the sufficient condition for an unbiased test no matter what pattern of confounding exists in the GWAS panel.

#### 3.3.1 Including 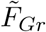 removes stratification bias

If we include 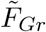 as a single fixed-effect covariate in the GWAS model, variation along 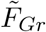 can no longer be used to estimate effect sizes. As a result 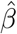 is uncorrelated with genotypes contrasts *r* under the null. If there is confounding along other shared axes of ancestry variation, the polygenic scores may still be biased along other axes, as

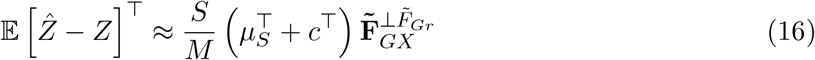

where

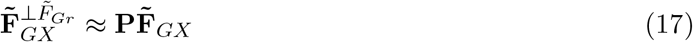

and 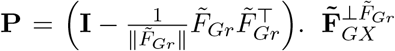 therefore captures cross panel relatedness along all axes of variation other than that specified by 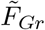. Controlling for variation aligned with 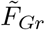 ensures that 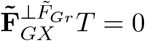, and it follows that

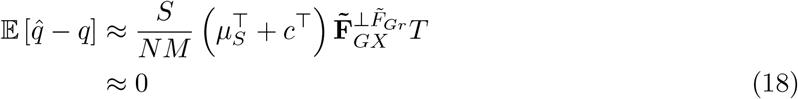

and the polygenic score association test is unbiased (see S5 and S6).

#### 3.3.2 Relationship between 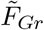 and PCA

A standard approach to controlling for population stratification in polygenic scores is to include the top *J* principal components of the GWAS panel genotype matrix as covariates in the GWAS, for some suitably large value of *J* [3]. In our model, how does this approach relate to including 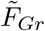 as a covariate in the GWAS?

As outlined in Section 2.1, **F**_*GG*_ contains the expected within panel relatedness for the individuals in the GWAS panel, the structure of which is determined by the demographic model. If we could take the eigendecomposition of **F**_*GG*_ directly, the resulting PCs are what we refer to as “population” PCs. The the number of population PCs that correspond to structure is entirely dependent on the population model. For example, below (section 3.4.1) we simulate under a 4 population sequential split model (Figure 1), in which case there are three population PCs that reflect real underlying structure. Later, (section 3.4.2) we simulate under a symmetric equilibrium migration model on a six-by-six lattice grid (Figure 3), in which case there are 35 population PCs reflecting underlying population structure. Including these population PCs as covariates in the GWAS would be sufficient to remove all ancestry-associated bias in effect size estimates and render the resulting polygenic scores uncorrelated with any axis of ancestry variation under the null hypothesis.

**Figure 1.**
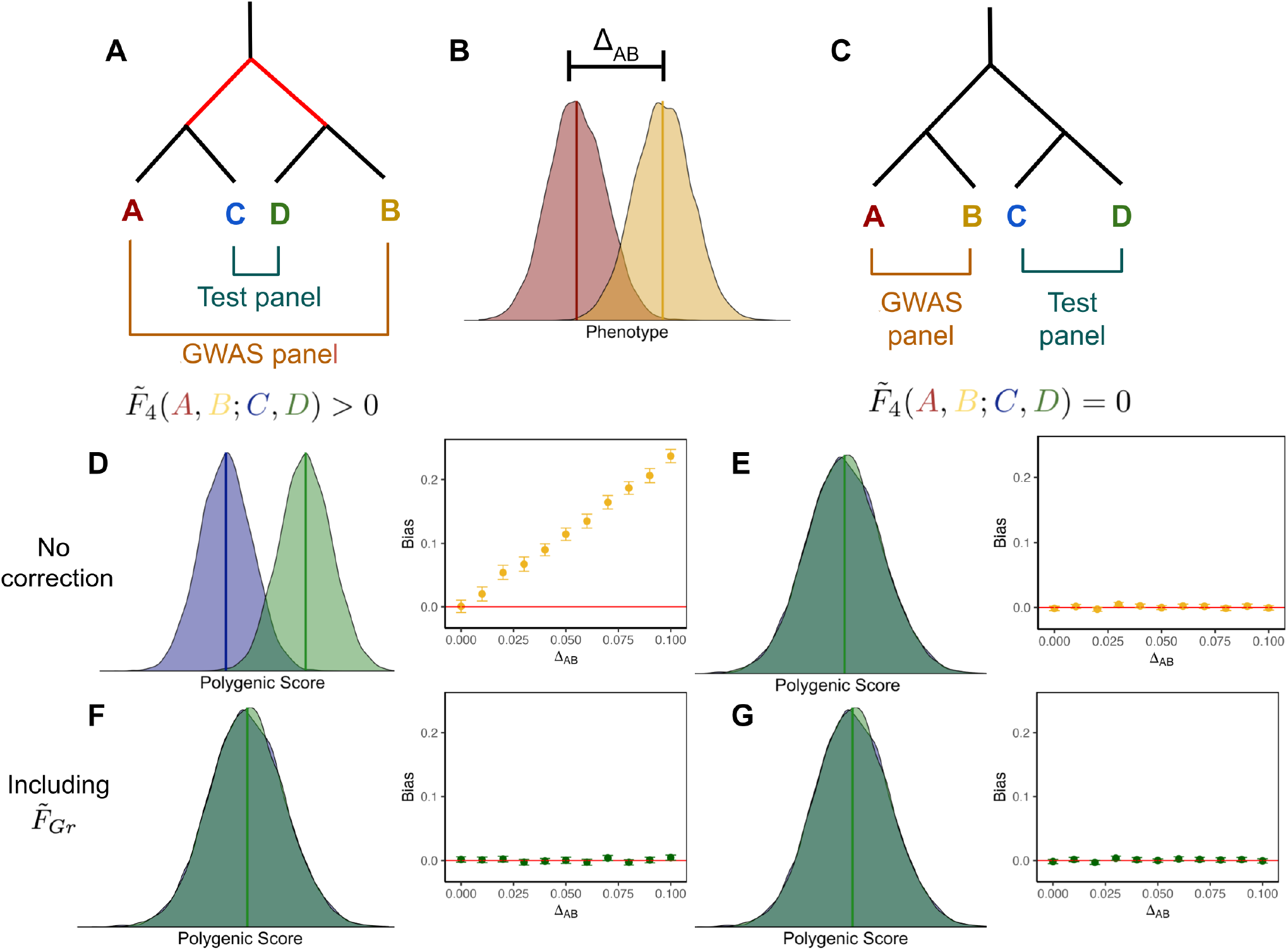
Schematic of two different panel configurations. The effect of stratification depends on the overlapping structure between the GWAS and test panels. (A, C) Two different topologies used to create the GWAS and test panels. (B) Stratification was modeled in the GWAS panel by drawing an individual’s phenotype *y* ∼ *N* (0, 1) and adding Δ_*AB*_ if they originated from population B. (D) When there is overlapping structure between GWAS and test panels, there is an expected mean difference between polygenic scores in populations C and D. Additionally, the bias in 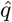 increases with the magnitude of stratification in the GWAS. (E) However, when there is no overlapping structure between panels, there is no expected difference in mean polygenic scores between C and D and 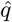 remains unbiased regardless of the magnitude of stratification. (F, G) Including 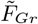 as a covariate in the GWAS controls for stratification, eliminating bias in 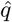 regardless of Δ_*AB*_ or the overlapping structure between GWAS and test panels.

**Figure 2.**
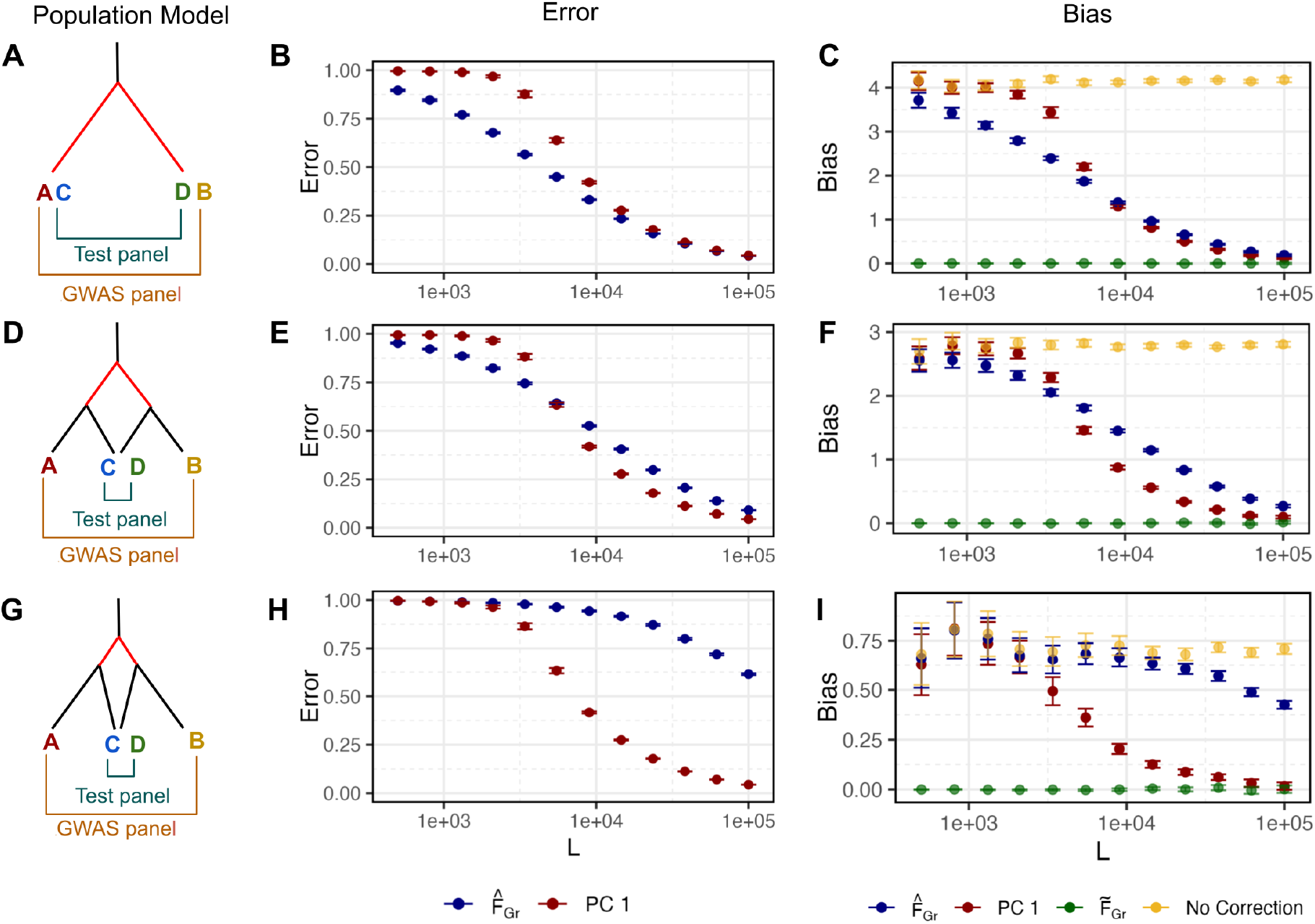
Error in estimators of 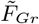 depends on the number of SNPs used to compute them. (A) We simulated a population model with a single split and sampled an equal proportion of individuals from each population to make a GWAS and test panel. (D,C) Here we simulated population models with two splits and sampled individuals in the overlapping structure configuration. (B, E, H) As 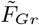 is known for these population models, we computed the error in *Û* _1_ and 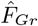 as estimators of 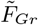 using eq. 27. For both estimators, error decreased as the number of SNPs increased. We hold the number of GWAS panel individuals constant at *M* = 1, 000 so as *L* increases the ratio of 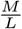 decreases. The error in *Û* _1_ does not depend on the population model as the depth of the deepest split is constant across models. Error in 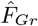 increases as overlap between panels decreases. (C, F, I) Bias in 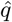 computed from using the estimators as covariates in the GWAS follows from the error in the estimators themselves.

**Figure 3.**
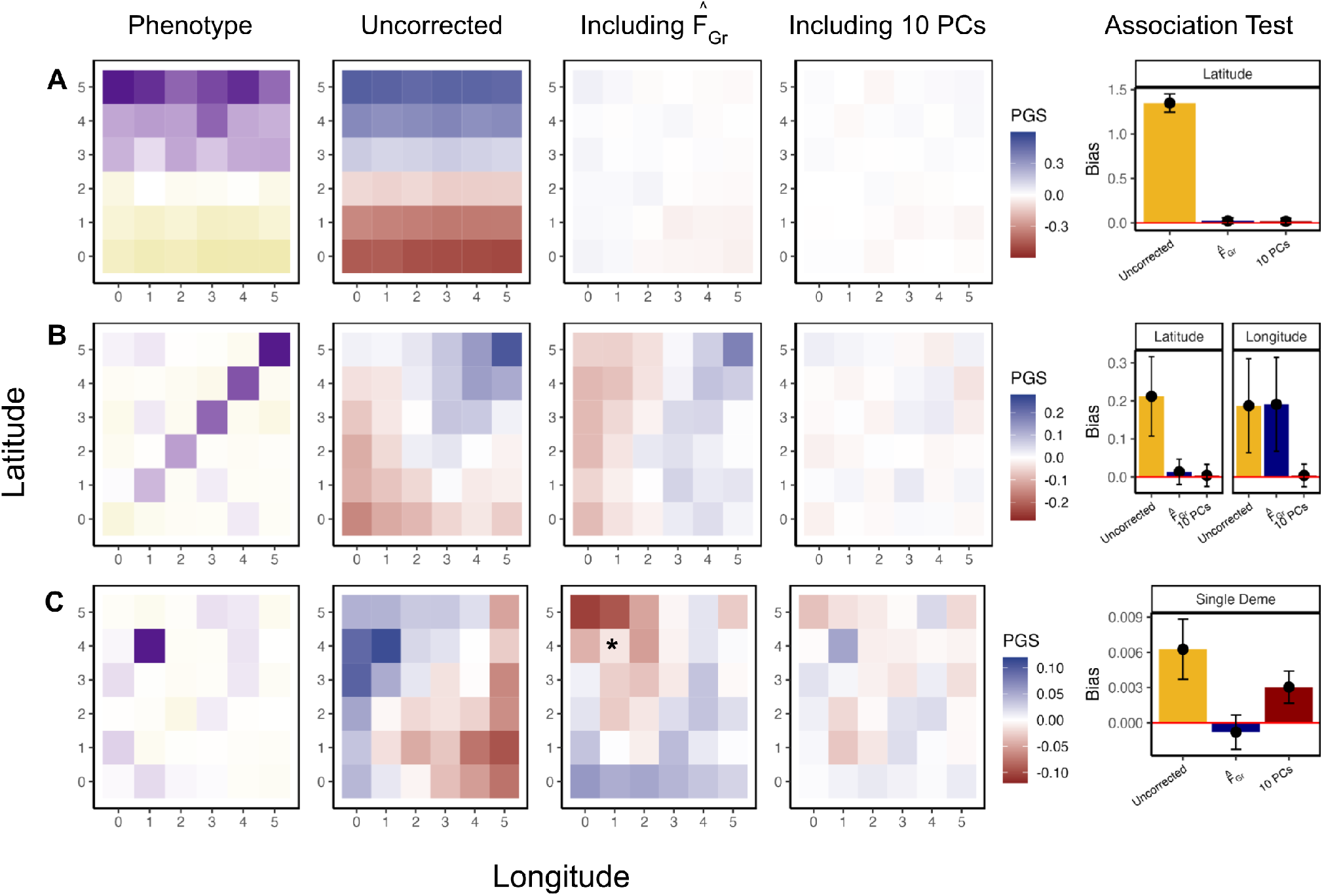
Stratification bias in more complex demographic scenarios. GWAS and test panel individuals were simulated using a stepping-stone model with continuous migration. In the GWAS panel, the phenotype is non-heritable and stratified along either latitude (A), the diagonal (B), or in a single deme (C). When effect sizes were estimated in a GWAS with no correction for stratification, polygenic scores constructed in the test panel recapitulate the spatial distribution of the confounder (second column). Including 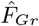 (test vector is latitude for A and B, belonging to * deme for C) in the GWAS model eliminates bias in polygenic scores along the test axis (third column) which is also reflected in the association test bias (fifth column). We also compare our approach to including the top 10 PCs (fourth column) which successfully protects the test in A and B but remains biased for C.

To see how the PCA correction approach works in the context of our theory, we can write 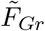 as a linear combination of GWAS panel population PCs,

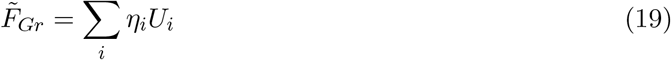

where *U*_*i*_ is the *i*^*th*^ PC of **F**_*GG*_ and the weights are given by 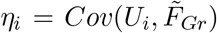. Estimating the marginal associations with 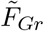 as a covariate can therefore be understood as fitting a model in which *all* population PCs are included as covariates, but the relative magnitude of the contributions from different PCs are fixed, and we estimate only a single slope that scales the contributions from all of the PCs jointly, i.e.

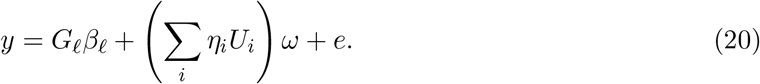

As a corollary, if we perform a polygenic score association test using GWAS effect size estimates in which the top *J* population PCs of **F**_*GG*_ are included as covariates, a sufficient condition for the included PCs to protect against bias from unmeasured confounders in a particular polygenic score association test is that 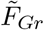 is captured by those *J* top PCs, i.e. that *η*_*i*_ ≈ 0 for *i > J*.

A second interpretation of the PC correction approach is that it operates on a hypothesis that the major axes of confounding in a given GWAS panel (i.e. *µ*_*S*_ and *c* in our notation) can be captured by the included PCs [45]. If this condition is met, effect size estimates are unbiased with respect to all axes of ancestry variation, whether they exist within a given test panel or not, and therefore any polygenic score association test that uses these effect size estimates will be unbiased with respect to ancestry as well. Combining this interpretation with results from above, population PCs should successfully eliminate bias in polygenic score association tests if the *J* PCs included in the GWAS either capture the confounding effects on the phenotype, eliminating all effect size bias, or if they capture 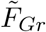, ensuring that effect size bias relevant to the test is removed.

#### 3.3.3 Controlling for bias in practice

Thus far we have shown the conditions under which including 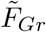 or the top J population PCs as fixed covariates removes stratification bias and leads to an unbiased association test. However, both 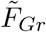 and *U* are theoretical quantities that depend on the population model, which we do not observe in practice. Instead, we must estimate these quantities, 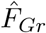 and *Û*, with error, from sample genotype data.

##### Sample principal components

The sample PCs, *Û*, can be computed by taking the eigendecomposition of the empirical genetic covariance matrix, or the singular value decomposition of the genotype matrix. Existing results from random matrix theory allow us to obtain some understanding of the accuracy of *Û* as an estimator of *U*. Specifically, in many GWASs the number of individuals in the GWAS panel, *M*, is roughly on the same order as the number of SNPs, *L*. In this setting, the accuracy of the sample eigenvector *Û* _*j*_ depends on the corresponding population eigenvalue (*λ*_*j*_) and the ratio of the number of individuals to the number of SNPs in the GWAS panel (^*M*^*/*_*L*_). As shown first by Patterson et al. (2006) in the context of genetics [46] (see also [47]), PCA exhibits a phase change behavior in which a given sample PC is only expected to align with the population PC if the corresponding population eigenvalue is greater than a threshold value of 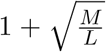. Below this threshold, the sample PC is orthogonal to the population PC.

However, even when the corresponding eigenvalue exceeds this threshold, the angle between the sample PC and the population PC may still be substantially less than one, particularly if the relevant eigenvalue does not far exceed the detection threshold [48, 49]. Specifically, the squared correlation between the population PC and the sample PC is approximately

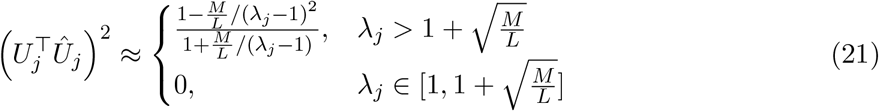

(see [48] for details). Thus even in cases where 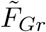 is fully captured by the top *J* population PCs, either of these two related phenomena may make it difficult to accurately approximate 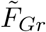 as a linear combination of the top *J* sample PCs, leading to a failure to fully account for stratification bias in polygenic score association tests.

##### Estimating 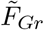 directly using test panel genotypes

Given this limitation of PCA, it’s natural to ask whether other estimators of 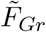 might perform better. One choice, suggested by our theoretical results, is a direct estimator that utilizes the relevant test panel genotype contrasts. Given the test panel genotype contrasts (*r*_*𝓁*_) and GWAS panel genotypes (*G*_*𝓁*_), we can obtain a direct estimator of 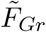 as

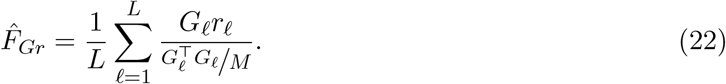

Then, if 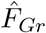 is a sufficiently accurate estimator of 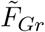, we should be able to render a given polygenic score association test unbiased by estimating marginal effects under the model

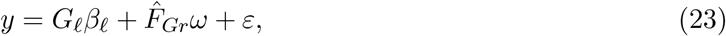

and ascertaining SNPs for inclusion in the polygenic scores via standard methods.

We can expect this method to be successful when the variance of the error component of 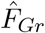 is small relative to the variance of the entries of 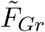. The variance of 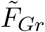 will be greater when the amount of overlap in population structure between the two panel along this specific axis is greater. We can think about the variance of the error component in terms of a linear model that tries to predict the GWAS panel genotypes using the test panel genotype contrasts. If we write 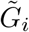· to denote the vectors of genotypes for GWAS individual *i* and 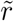 for the test panel genotype contrasts, each standardized by the variance in the GWAS panel, then we can fit the linear model

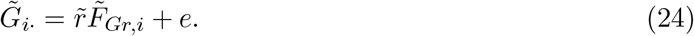

The regression coefficient estimate from the fitted model is then the *i*^*th*^ entry in our population structure estimator, 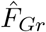. The error in 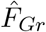 therefore behaves like the error in a typical regression coefficient, and should be minimized when the number of SNPs included, *L*, is large, and when the test panel sample size, *N*, is large, so that the 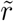 are well estimated.

This approach proposes to use the test panel genotype data twice: once when controlling for stratification in the GWAS panel, and a second time when testing for an association between the polygenic scores and the test vector. One concern is that this procedure might remove the signal we are trying to detect. In supplemental section S7.1 we show that while this is true for naive applications, the effect will be small so long as the number of SNPs used to compute the correction is large relative to the number included in the polygenic score (i.e *S ≪ L*). Notably, controlling for sample PCs of the GWAS panel genotype matrix will induce a similar effect if the sample PCs capture 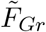. We confirm via simulations (see supplemental section S7.2, and Figure S3) that downward bias in 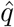 when including 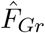 or sample PCs is minimal when *S ≪ L*. Further concern about downward biases in applications could likely be ameliorated via the “leave one chromosome out” scheme commonly implemented in the context of linear mixed models [50, 5] or via iterative approaches that first aim to ascertain SNPs using a genome-wide estimate of 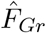 before re-estimating effects using an estimate of 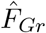 computed from sites not in strong LD with any of the ascertained sites.

### 3.4 Applications

In this section, using theory, simulations and an application to real data, we consider a number of concrete examples with varying degrees of alignment between the axis of stratification and axis of population structure relevant to the polygenic score association test, demonstrating how these biases play out in practice, and how well PCs and 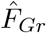 capture bias in different circumstances.

#### 3.4.1 Toy Model

##### Stratification bias depends on 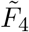 (A, B; C, D)

We first consider a toy model with four populations (labeled A, B, C and D), which are related to one another by an evenly balanced population phylogeny (Figure 1). The GWAS panel is composed of an equal mixture of individuals from populations A and B, and we test for a difference in mean polygenic score between populations C and D under two different topologies, one where A and C are sister to one another (Figure 1A), and another where A and B are sister (Figure 1C).

For simplicity, we consider a purely environmental phenotype (i.e. *h*^2^ = 0) with a difference in mean between populations A and B equal to Δ_*AB*_ (Figure 1B). Following from eq. 7, the marginal effect size estimate for site 𝓁 is

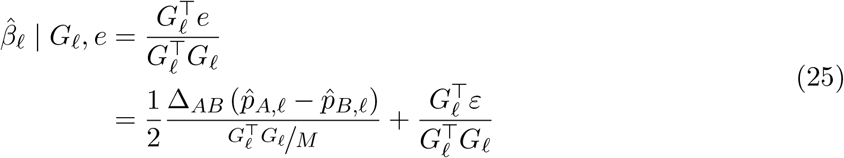

where 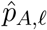 and 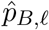 are the observed sample allele frequencies for population A and B at site 𝓁 (see also equation 2.3 in the supplement of [27]).

Then, using these effect sizes to test for a difference in mean polygenic score between populations C and D, the bias in our association test statistic is,

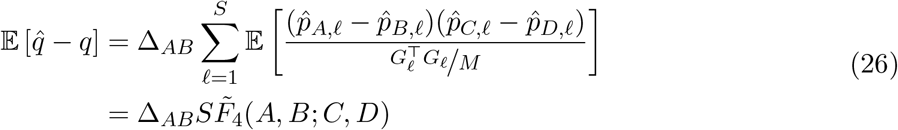

where 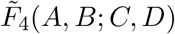 is a version of Patterson’s *F*_4_ statistic [51, 52], standardized by the genotypic variance in the GWAS panel, which measures the amount of genetic drift common to populations A and B that is also shared by populations C and D. Writing the bias in terms of this modified *F*_4_ statistic helps illustrate the role of cross panel population structure in driving stratification bias in polygenic scores. The effect estimate at site 𝓁 is a linear function of 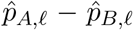, so the test will be biased if 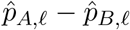 is correlated with 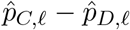. This is true for the demographic model in Figure 1A, where shared drift on the internal branch generates such a correlation, yielding a positive value for 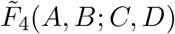, but not for the model in Figure 1C, where there is no shared internal branch and 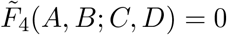.

To test this prediction, we simulated 100 replicates of four populations related by this topology. In the GWAS panel populations we simulated purely environmental phenotypes with a difference in mean phenotype (as outlined above), conducted a GWAS, ascertained SNPs, and then used these SNPs to construct polygenic scores and compute 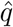 in the test panel. The results are consistent with our theoretical expectations: the test statistic is biased for the topology with 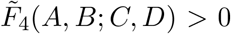 (Figure 1D), but unbiased when 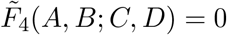 (Figure 1E).

Given the population model, 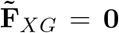 for the unconfounded topology, making 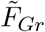 a vector of zeros. Therefore, re-running the GWAS including 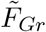 does not change the outcome of the already unbiased test (Figure 1G). For the confounded topology, the structure in 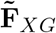 reflects the deepest split in the phylogeny and is aligned with *T*. 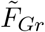 is therefore an indicator of which GWAS panel individuals are on which side of the deepest split and including it as a covariate in the GWAS eliminates bias for the confounded topology (Figure 1F).

##### Quantifying error in estimators of 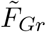

As we outlined above, in practice, 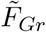 cannot be observed directly, and must be estimated with error from the data. To illustrate the impact of this estimation error on the performance of both estimators in a simple, well understood case, we performed simulations using three different versions of our toy model in which we vary the amount of overlap in population structure between the test and GWAS panels. Specifically, given that 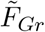 is known in this toy model, we can compute the error in either estimator as one minus the squared correlation between 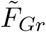 and the corresponding estimator. We take all of these vectors to be standardized, so this is simply

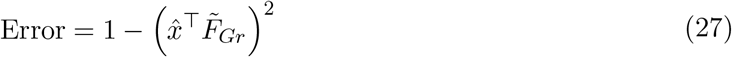

where 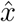 represents the appropriate estimator.

For each simulation, we estimated 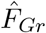 as in eq. 22, using *L* genome-wide SNPs with a frequency of greater than 1% in both the test and GWAS panels. For PCA, we computed sample PCs via singular value decomposition of the genotype matrix using the same set of SNPs that were used to compute 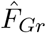, and we then take *Û* _1_ (i.e. the first sample PC) as the PCA based estimator of 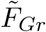 [42]. In all of these simulations, we hold the GWAS and test panel sample sizes constant at *N, M* = 1, 000 and varied the number of SNPs (*L*) as a way to vary the accuracy of the estimators. We simulated 100 replicates for each topology, and plot the resulting averages across these replicates in Figure 2.

First, we simulated a scenario of complete overlap, in which there is a single population split and individuals in both the GWAS and test panels are independently drawn as 50:50 mixtures of the two population on either side of the split (Figure 2A). When the GWAS sample size (*M*) is on the same order as the number of SNPs (*L*), the direct estimator 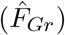 has a smaller error than the first PC (*Û* _1_) (Figure 2B), and as a consequence reduces the bias by a larger amount (Figure 2C). Intuitively, the direct estimator singles out the relevant axis of population structure because we have identified it ourselves in the test panel, whereas PCA has to find this axis “on its own” in the high dimension GWAS panel genotype data, and thus pays an additional cost. In contrast, when *M ≪ L* so that ^*M*^*/*_*L*_ ≈ 0, PCA no longer has to pay this additional cost, and its performance improves to match that of the direct estimator.

We next simulated under the same toy model of partial overlap in population structure between test and GWAS panels that we considered above in Figure 1 (Figure 2D). This results in an increase in the error of the direct estimator relative to the complete overlap case because the genotype contrasts measured in the test panel are less informative about the relevant axis of structure in the GWAS panel. In contrast, the error in *Û* _1_ is unchanged, as the amount of structure in the GWAS panel is the same as inFigure 2A. Notably, in this case the direct estimator still outperforms PCA when ^*M*^*/*_*L*_ *>* 0, but PCA performs better when ^*M*^*/*_*L*_ ≈ 0.

Finally, in Figure 2G we reduced the overlap in population structure even further, which leads PCA to uniformly outperform the direct estimator, even in the ^*M*^*/*_*L*_ *>* 0 regime. Intuitively, because the overlap in population structure is so small, the direct estimator requires a very large number of SNPs to produce an accurate estimate. We also note that in general across all of these simulations, while the magnitude of the reduction in bias closely tracks the error in the estimator of population structure, the reduction is slightly larger than expected for *Û* _1_ (Figure S1).

#### 3.4.2 Grid Simulations

To further explore stratification bias in more complex scenarios, we conducted another set of coalescent simulations under a symmetric two-way migration model on a six-by-six lattice grid, building off of a framework developed by Zaidi and Mathieson (2020) [53]. We sampled an equal number of individuals per deme to comprise both the GWAS and test panels, with total sample sizes *N, M* = 1, 440. We then simulated several different distributions of purely environmental phenotypes across the GWAS panel individuals. We considered three different scenarios for the distribution of phenotypes. For each scenario, we estimated effect sizes, ascertained associated sites, and tested for an association between polygenic score and latitude, longitude, or membership in the single confounded deme, depending on the example. In these simulations 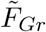 is unknown and so we compared 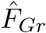 and the top 10 sample PCs as estimators of 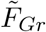, using the same set of *L* = 20, 000 SNPs that are found at a frequency greater than 1% in both panels for both estimators.

For the first example, the confounder, *c*, is a linear function of an individual’s position on the latitudinal axis (Figure 3A). When we estimated effect sizes with no correction for population structure, the spatial distribution of the resulting polygenic scores reflected the distribution of the environmental confounder. Consequently, an association test using latitude as the test vector is biased. However, including 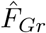 or the top 10 sample PCs as covariates in the GWAS model is sufficient to ensure that effect sizes that are unbiased with respect to the latitudinal genotype contrasts in the test panel, so the resulting association test is unbiased.

In the second example, we simulated confounding along the diagonal, resulting in uncorrected polygenic scores that are correlated with both latitude and longitude in the test panel and an association test that is biased along both axes (Figure 3B). When we computed 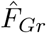 using latitude as the test vector, the resulting effect sizes are uncorrelated with latitudinal genotype contrasts, but remain susceptible to bias along other axes (e.g. longitude). This example highlights the targeted nature of this approach, as using effect sizes from a GWAS including 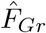 does not remove all bias, but does make the association test using those effect sizes for the pre-specified test vector unbiased (when 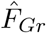 is well estimated). Including 10 sample PCs protects both the latitudinal and longitudinal association tests.

In the third example, we simulated an increased environmental effect in a single deme, a scenario which induces a more complex spatial pattern in the uncorrected polygenic scores (Figure 3C), and which previous work has shown to be difficult to correct for with standard tools [54, 53]. We then took the test vector to be an indicator for whether the test panel individuals were sampled from the deme with the environmental effect or not, and compute 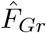 using these contrasts. In this scenario, including 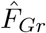 as a covariate in the GWAS results in an unbiased test statistic. In contrast, the top ten sample PCs did not.

##### Quantifying error in population structure estimators

Next, we wanted to better understand the role of error in our population structure estimators plays in these simulations. In contrast to the four population toy model, it is not straightforward to compute 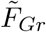 given our underlying demographic model, particularly for the case of testing a single deme against all others. As a result, we cannot directly measure the error in 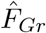 or sample PCs as estimators of 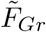. Instead we use the fact that under this demographic model individuals within a deme are exchangeable, and therefore have the same values of both 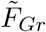 and population PCs. This allows us to estimate the error in 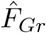 by computing the fraction of the total variance in 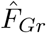 that can be attributed to variance of individual values within demes, and to variance of deme means across replicates (see Section 5.6.1). For the PCs the relationship between the order of the underlying population PCs and the order of the sample PCs may differ across replicates due to the noisiness of the sample PCs so it is not obvious how to compute the variance of the deme means across replicates. We therefore use only the within deme variances, so our estimates of the error for the PCs are technically estimates of a lower bound on the error (see Section 5.6.2). However, we note that for our estimation of the error in 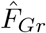, we found that the variance within demes was by far the larger contributor, so we expect this to be a relatively tight bound. We then vary the number of SNPs used to compute our estimators of population structure from *L* = 20, 000 down to *L* = 2, 000, and observe how differences in the estimated error of our population structure estimators translate to differences in the amount of bias in the polygenic score association test statistic.

In Figure 3A and Figure 3B, 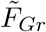 corresponds to latitude, so we expect it to be captured by the top two population PCs [55]. For *L* = 20, 000 (the number of SNPs used in Figure 3), we estimated the lower bound on the error in sample PCs 1 and 2 to be 0.011. Across the range of *L* values we tested, the estimated bound was no greater than 0.053 (Figure 4A) and including 10 PCs consistently removes bias in 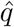 (Figure 4B). Similarly, we estimated the error in 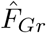 for latitude to be 0.012 when *L* = 20, 000 with a maximum of 0.059 when *L* = 2, 000. Although these estimates are nearly identical to the values we observe for the first two PCs, the bias in 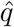 is slightly higher (Figure 4B). We observed a similar result in the 4 population toy model (Figure S1), so this may be the same phenomenon, or it may be that PCs 3-10 are capturing some of the residual latitudinal signal that is not captured by the first two.

**Figure 4.**
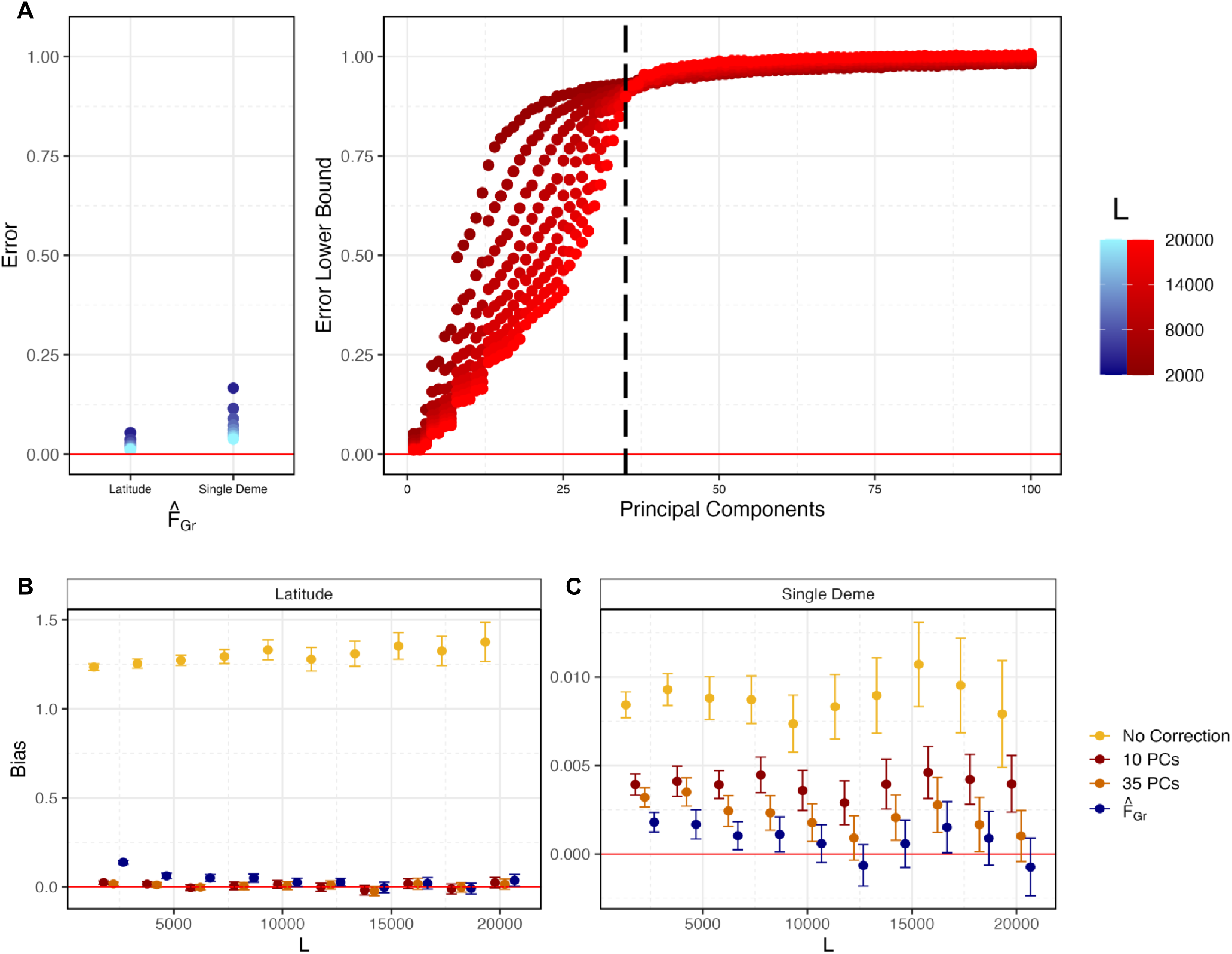
Quantifying error in estimates of 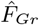 and sample PCs for the six-by-six stepping stone demographic model. (A) Given the stepping stone demographic model used in Figure 3, individuals within a deme are exchangeable and have the same 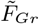 and population PC value. Therefore we used variation within demes to estimate the error in 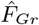 and a lower bound for the error in sample PCs (see Section 5.6.1 and Section 5.6.2 for details) for different values of *L* (we hold *M* = 1, 400). The dashed vertical line indicates PC 35, the last population PC we expect to capture real structure. (B) When latitude is the test vector, both sample PCs and 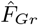 are well estimated and bias in 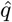 is reduced. (C) When a single deme indicator variable is the test vector, higher PCs are needed to capture 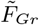. These sample PCs are not well estimated and residual bias remains when 35 PCs are used for most values of *L*.

Next, we explored the role of error in our population structure estimators for the more difficult single deme test/confounder case (Figure 3C). We again computed the error in 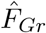 as we vary *L*, with estimates ranging from 0.04 to 0.18 as *L* decreases (Figure 4A). For larger values of *L*, the error was small enough that confidence intervals on the bias overlapped zero, but this was not true when we reduced *L* so that the error was larger (Figure 4C). Above, with *L* = 20, 000, we found that 10 PCs were not sufficient to remove the bias. This could either be because 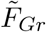 is not captured by the top 10 population PCs *or* it could be that 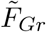 can be captured by 10 population PCs, but the sample PCs are too noisy as estimates of the population PCs. Given that there are 36 demes in our simulations and that individuals within demes are exchangeable, only the top 35 population PCs capture real population structure, while the rest correspond to sampling variance. As a result, if the sample PCs are sufficiently well estimated, then only 35 should be required to remove the bias. In practice, we find that using 35 PCs for larger values of *L*, the bias is closer to zero than it is with 10 PCs, but the confidence intervals still to do not always overlap zero, and the bias is generally greater than it is when we use our direct estimator, 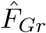 (Figure 4C). As expected, the performance with 35 sample PCs decreases further with an increase in the error, but is always intermediate between 10 PCs and 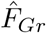. All of this is consistent with the observation that the error in the higer sample PCs (i.e. 11-35), is extremely high across the range of *L* values we explored (Figure 4A).

##### PCs succeed by capturing structure relevant to the test, not the confounder

Finally, to the extent that the PCs did succeed in removing bias in our simulations, we wanted to understand whether it was because they successfully captured the confounder, or because they captured the relevant axis of structure for the test (see section 3.3.2). To this end, for each of the three grid scenarios in the *L* = 20, 000 case, we computed the cumulative proportion of variance in the confounder, *c*, that could be explained by the first *J* sample PCs, for *J* up to 100 (Figure 5). We found that while the confounding axis was well captured by sample PCs 1 and 2 for latitude (Figure 5A), it was not well captured by the top 10, 35, or indeed 100 PCs for the diagonal (Figure 5C) or single deme confounders (Figure 5E). In contrast, if we take our estimator, 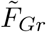, as a proxy for 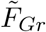, we find that the PCs explain a considerably higher fraction of the variance. For the first two cases, the test axis is latitude, so this is unsurprising. However, this is true even for the single deme case, and results from the fact that relatedness among adjacent demes leads in a smoothing effect (Figure S2), which makes 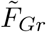 easier for the PCs to capture.

**Figure 5.**
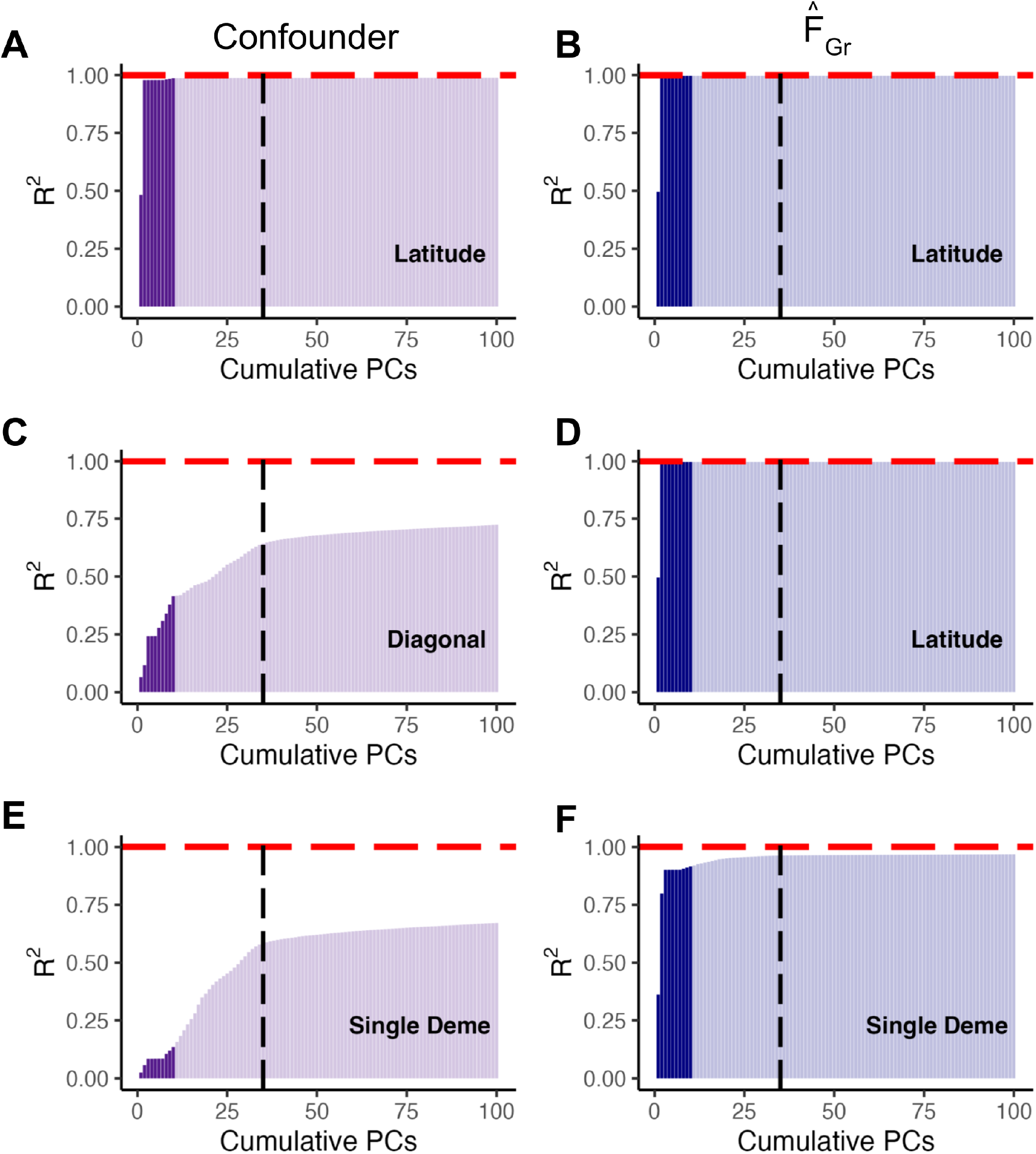
Different patterns of confounding and 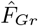 are captured by different GWAS panel sample PCs. For the three possible combinations of confounding and polygenic score association tests in Figure 3, we plot the variance in either the confounder or 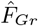 explained by cumulative GWAS panel sample PCs, with the top 10 PCs highlighted in a darker color. As 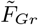 is unknown for this model, we estimated the error in 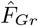 as 0.011 and 0.04 for latitude and the single deme, respectively, and therefore assume it is a decent proxy for 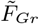. In (A) both the confounder and 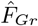 (and therefore 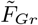) represent variation along latitude and are well captured by the first two PCs. For (B) the confounder varies along the diagonal and these individual deme level differences are not well captured by top sample PCs. In contrast, the test vector is still latitude and 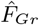 is again well captured by PCs 1 and 2. Finally, in (C), both the confounder and the test vector represent membership in a single deme and therefore not as well captured by top sample PCs.

## 4 Discussion

Interpreting patterns in the distribution of polygenic scores is difficult, especially when confounding cannot be ruled out. Because most well-powered GWAS are conducted on population samples where the relationship between genetic background, ancestry, and the environment is not well controlled, stratification bias remains a significant concern [32, 33, 40, 56]. Here, we characterize patterns of stratification bias in the distribution of polygenic scores as a function of the expected genetic similarity between GWAS and test panels. For any given polygenic score association test axis, the amount of bias in the association test statistic depends on the strength of stratification along exactly one axis of population structure in the GWAS panel 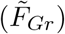.

The ability to conduct a given polygenic score association test in an unbiased manner therefore depends on the accuracy with which we can model 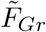 via co-variates included in the GWAS. For the standard PCA based approach the inconsistency of the sample PCs as estimators of population structure is therefore a plausible explanation for the signatures of residual stratification bias that have been reported across many GWAS datasets [32, 33, 40], though such signals might also arise simply from not including enough PCs, even if they are well estimated. The inconsistency of the sample PCs as estimators is a well known result in random matrix theory [47, 48], and we are not the first to notice the connection to stratification bias in GWAS and polygenic scores [49], but the phenomenon is not widely acknowledged in the GWAS literature.

In light of these issues, we proposed a direct estimator of the target axis of population structure using the test panel genotype data, and show that under optimal conditions of complete overlap in structure between panels and a large sample size in the test panel (Figure 2A and Figure 4C) this estimator outperforms, or at least equals, the standard PCA based estimator. A limitation this direct approach is that the performance relative to PCA degrades as the amount of overlap in structure between the two panels decreases (Figure 2B and Figure 2C). As a result it is best suited to cases where the GWAS cohort and test panels are drawn from the same sample, thus ensuring a high overlap in structure between panels. We also expect this method to perform best when the test panel is large relative to the amount of variance explained by the test vector, so that the relevant genotype contrasts, *r*, are well-estimated.

Several recent papers have proposed alternative methods for improved control of population structure in GWAS and polygenic scores. Proposals include using 1) PCs of rare variants (as opposed to common variants) [53], 2) PCs of external reference datasets in addition to the PCs of the GWAS panel [57], 3) or local ancestry assignments (in lieu of global linear estimators) [58]. Our results highlight the importance of developing tools to more robustly estimate the error in population structure estimates [59], and it would be interesting to understand the merits of these alternative methods through this lens. Ideally, future methods development might allow each set of GWAS summary statistics to be accompanied by statistics summarizing the accuracy of the population structure estimates used to control for stratification. These estimates could then be used in downstream analyses to provide quantitative statements about the extent to which a particular polygenic score association test is or is not protected from stratification bias. We also note that tests for association between polygenic scores and axes of ancestry variation are closely related to bivariate LD score regression as applied to a combination of effect estimates for one trait and frequency/genotype contrasts from an independent dataset [60, 19, 32]. Previous work in the context of polygenic selection tests raised concerns about spurious inflation of the LD score slope due to background selection [32]. It would be interesting to revisit this issue more fully in light of our present results.

There are several elements of our model that differ from reality. It is worth highlighting what these are, and what their effects are. For example, our model ignores linkage among sites and assumes that we use marginal effects, rather than jointly estimated effects, to construct our polygenic scores. Firstly, linkage among sites does not change the fundamental point that controlling for 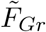 is sufficient to render the effect size estimates uncorrelated with the test panel genotype contrasts under the null. This is true whether effects are estimated marginally or jointly. However, in practice, we would still prefer to estimate effects jointly, for at least two reasons. The first is simply because doing so increases the accuracy of the polygenic score, which will increase our power. The second is because, in the presence of residual stratification (e.g. if our estimator, 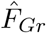, has high error), polygenic scores constructed with jointly estimated effects should be less biased than those constructed using marginal effects. This is because, when effect sizes are estimated marginally, each site experiences the entirety of the stratification effect, and therefore gets a “full dose” of it. The stratification effect is then being added into the polygenic score multiple times across SNPs. This is why we find the bias in the polygenic score association test statistic to be proportional to the the number of loci included in the polygenic score. In contrast, if effects were estimated jointly, the stratification effect will be spread out more evenly across sites, and so we would expect the effect on the polygenic score to be less extreme, but not eliminated.

Another issue is that, throughout our simulations we often estimate effect sizes while attempting to control for stratification *only* along the target axis of the test. We do this to highlight our main point that controlling for the target axis is sufficient to render the association test unbiased, but readily acknowledge that it does not deal with all of the negative consequences of stratification bias. For example, bias along other axis will function as additional noise in the process of ascertaining SNPs, and in the polygenic scores themselves, which would be expected to reduce power. Therefore, it is still desirable to include top PCs or use a LMM alongside 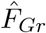, even in the case where 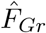 is well estimated.

We also wish to emphasize that our results are relevant for a broader set of analyses than those explicitly covered by our model. For example, with a slight shift in perspective, our model is applicable to studies that use GWAS summary statistics together with coalescent methods to test for signals of directional polygenic selection [19, 23, 24, 61]. The key to this is to recognize such methods use patterns of haplotype variation to estimate genotype contrasts between the sampled present day individuals and a set of unobserved ancestors, and then ask whether these estimated genotype contrasts correlate with effect size estimates for a trait of interest. Thus, within such an analysis there also exists an 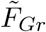 that describes the extent to which individuals in the GWAS panel are more closely related to the present day sample or the hypothetical ancestors. For both the coalescent approaches, as well as methods relying on direct comparison of polygenic scores, both the evolutionary hypothesis being tested and the degree of susceptibility to bias follow directly from the set of genotype contrasts used in the test. Some prior work has suggested that certain coalescent methods of testing for polygenic selection are more robust to stratification bias than others [24, 61], but our results show that this cannot be true: two different methods that test the same evolutionary hypothesis using the same set of estimated effect sizes necessarily have the same susceptibility to stratification bias. If there *are* differences in robustness to stratification bias among methods, then this must come either from changing the evolutionary hypothesis being tested or from overall differences in the statistical power of the methods.

Finally, we note that even if 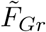 is known exactly the interpretation of the results of polygenic score association tests is limited by the many assumptions that must be made in any polygenic score analysis [62]. For example, these analyses use effect sizes estimated in a one set of genetic and environmental background, and there is no guarantee that the effects will be the same in other backgrounds. Effect size heterogeneity can cause many difficulties with the interpretation of positive associations between polygenic scores and axes of population structure (as several papers have noted [62, 13, 63]). Another difficulty with interpretation arises from allelic turnover [38] and differences in tagging across populations, as a given polygenic score will have less power to detect differences between populations that are genetically more distant from the GWAS panel, and this can lead to a biased picture of how selection has actually affected the trait across populations [39]. However, none of these phenomena are expected to generate false signals of directional selection where none exists. This is because the fact that the effect size might vary across populations has no impact on the correlation between the effect size measured in only one of the populations and patterns of allele frequency differentiation among populations. One subtle caveat to this claim is that certain forms of directional interaction effects (e.g. directional dominance) could in principle create correlations between the direction of recent allele frequency change on the lineage leading to the GWAS panel individuals and the average effect as estimated under an additivity assumption, and this *would* violate the null model. However, there is little evidence for substantial interaction variance among common variants in human complex traits, so this is unlikely to be an issue in practice.

Moving beyond the specific issue of associations between polygenic scores and population structure axes, we note that GWAS can also be impacted by other forms of genetic confounding beyond the simple associations between ancestry and genetic background that we consider here, include dynastic effects, assortative mating, and stabilizing selection [64]. Therefore, while our results provide a pathway to a more rigorous approach for protecting against stratification bias in polygenic score association tests, addressing a known problem in their implementation, continued care in the interpretation of polygenic score analyses is always warranted.

## 5 Materials and Methods

### 5.1 Simulating genotypes

We used *msprime* [65] to simulate genotypes under different models with 100 replicates per model. The first model, shown in Figure 1, has two population splits, 200 and 100 generations in past, for a total of 4 present day populations. We fix the population size for all present and past populations to 10,000 diploid individuals. We then sample 5,000 individuals per population and create two configurations of GWAS and test panels (*N, M* = 10, 000) based on the diagrams in Figure 1A and Figure 1C. For every model replicate we simulate a large number of independent sites and downsample to *L* = 10, 000 SNPs with MAF *>* 0.01 in both GWAS and test panels. We use these genotype simulations for Figure 1 and Figure S3. When the populations in the GWAS and test panel are non-sister (i.e Figure 1A) the average within panel *F*_*ST*_ [66] was 0.01, whereas in the configuration in Figure 1C the average *F*_*ST*_ was 0.005.

For Figure 2 we use the same model setup but adjust the split times to 12/0, 12/4, and 12/10 generations in the past for population models A, B, and C, respectively. The average *F*_*ST*_ for the overlapping structure scenario is approximately 0.0006. To reduce computational burden, we scale down the sample size to 1,000 individuals per panel (500 per population). We simulate large number of independent SNPs and down-sample to *L* sites (MAF *>* 0.01 in both panels) which we vary from 500 to 100,000.

For Figure 3 we use a model, modified from [53], that is a 6 × 6 stepping stone model where structure extends infinitely far back with a symmetric migration rate of *m* = 0.01. We fix the effective population size to 1,000 diploid individuals and sample 80 individuals per deme which we split equally into GWAS and test panels (*N, M* = 1, 440). As above, we simulate large numbers independent SNPs and down-sample to *L* = 20, 000 SNPs with MAF *>* 0.01 in both panels.

### 5.2 Simulating phenotypes

To study the effect of environmental stratification on association tests, we first simulated non-genetic phenotypes for an individual *i* in the GWAS panel as *y*_*i*_ ∼ *N* (0, 1). In our discrete 4 population models we then generate a phenotypic difference between populations by adding Δ_*AB*_ to *y*_*i*_ for individuals in population B. For Figure 1 we vary Δ_*AB*_ from 0 to 0.1 standard deviations. In order to compare across models and values of 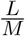 in Figure 2 we compute 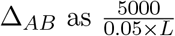.

In our grid simulations we generated three different phenotypic gradients where the largest phenotypic shift was always equal to Δ. To generate a latitudinal gradient (Figure 3A) we added 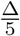 to *y*_*i*_ for individuals in row 1, 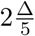 for individuals in row 2, etc. For Figure 3B we generated a gradient along the diagonal by adding 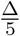 to the phenotype for individuals in deme (1,1), 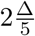 for individuals in deme (2,2), etc. For Figure 3C we shifted the phenotype of individuals in deme (1,4) by Δ. For all grid simulations in Figure 3 we set Δ = 0.2. In order to compare across values of *L* in Figure 4 we compute Δ as 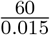

To study the effect of controlling for stratification in cases where there is a true signal of association between polygenic scores and the test vector (Figure S3), we used our 4 population demographic model and followed the protocol outlined in [53] to simulate a neutral trait with *h*^2^ = 0.3. We first randomly select 300 variants to be causal and sample their effect sizes from 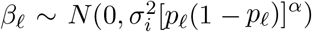, where 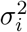 is a frequency independent scale of the variance in effect sizes, *p*_*𝓁*_ is allele frequency in the GWAS panel, and *α* is a scaling factor controlling the relationship between allele frequency and effect size. We set *α* = −0.4 and 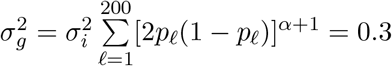.

To simulate a signal of true difference in polygenic score in the test panel, we calculate the frequency difference *p*_*D,𝓁*_ − *p*_*C,𝓁*_ at all 300 causal sites in the test panel and flip the sign of the effect sizes in the GWAS panel such that *p*_*D*_ − *p*_*C*_ *>* 0 and *β*_*𝓁*_ *>* 0 with probability *θ. θ* therefore controls the strength of the association with *θ* = 0.5 representing no expected association and *θ* = 1 representing the most extreme case where trait increasing alleles are always at a higher frequency in population D. We use *θ* ranging from 0.5 − 0.62. We then draw the environmental component of the phenotype *e*_*i,k*_ ∼ *N* (0, 1 − *h*^2^) and generate an environmental confounder by adding Δ_*AB*_ ∈ {−0.1, 0, 0.1} to *e*_*i,k*_ for individuals in population B.

### 5.3 Computing covariates

For each polygenic score association test we computed 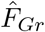. We first construct *T* as either population ID, latitude or the single deme of interest, depending on the simulation. Given this test vector, we compute *r* = **X**^*T*^*T* using the plink2 [67] function `--glm`. Finally we compute 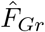 (see eq. 22) using `--sscore` in plink2, taking care to standardize by the variance in the GWAS panel genotypes. Additionally we used plink2 [67] `--pca` or `--pca approx` to compute sample PCs from the GWAS panel genotype matrix.

### 5.4 GWAS

For each set of phenotypes, we carried out three separate marginal association GWASs using the regression equations below,

1. *y= β* _*𝓁*_*G*_*𝓁*_ + ϵ
2. 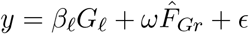
3. *y= β* _*𝓁*_*G*_*𝓁*_ + *ω*_1_ *Û*_1_ + … + *ω*_*j*_ *Û*_*j*_ + *ϵ*

Additionally, we conducted a fourth GWAS, 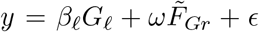, for the discrete 4 population model where 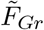 is known. All GWASs were done using the plink2 [67] function `--glm`.

We then ascertain *S* SNPs based on minimum p-value for inclusion in the polygenic score. For Figure 1 and Figure 3 we set *S* = 300. In order to compare across values of 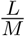 in Figure 2 and Figure 4, we set *S* = 0.05 × *L* and *S* = 0.015 × *L*, respectively. For Figure S3 we use use estimated effect sizes at the 300 causal sites rather than ascertaining based on p-value.

### 5.5 Polygenic Score Association Test

We construct polygenic scores for the individuals in the test panel as 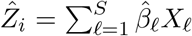 where 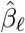 is the estimated effect size from the joint model and *X*_*𝓁*_ is the mean centered genotype value for the 𝓁^*th*^ variant.

For each replicate we then compute the test statistic 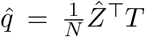 by multiplying the vector of polygenic scores for individuals in the test panel by the test vector. Finally we compute the bias in 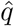 across each set of 100 replicates as 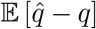.

### 5.6 Estimating the error in our population structure estimators for the grid model

#### 5.6.1 Direct estimator

Consider that the value of 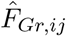, the entry of 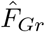 for the *i*^*th*^ individual in the *j*^*th*^ deme, can be decomposed as

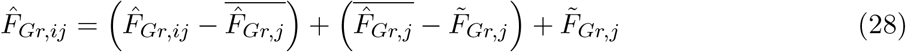

where 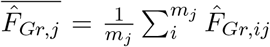 is the empirical average of 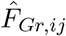 within deme *j* (*m*_*j*_ is the number of individuals in deme *j*), and 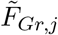 is the entry of the true population structure axis 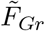, for all individuals in deme *j*. Individuals within demes are exchangeable in our model, so the deviations 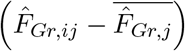 and 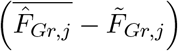 both represent sources of error in our estimator. The fraction of variance in 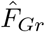 that is attributable to error is therefore

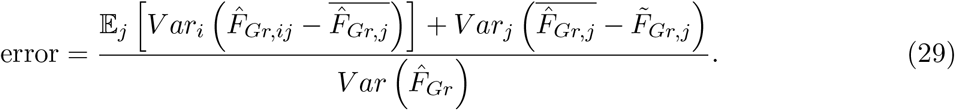

We can estimate 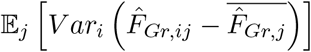 as

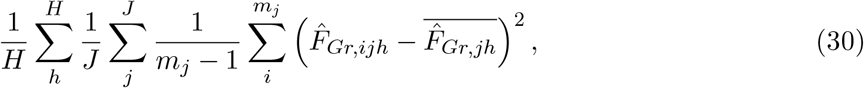

where *h* indexes replicate simulations and *H* is the total number of replicates (*H* = 100 in our case), *J* gives the total number of demes (36 in our case), *m*_*j*_ is the number of individuals in deme *j*, and

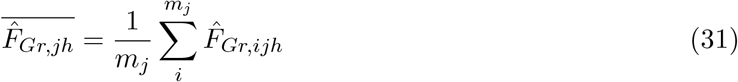

is the empirical mean entry for deme *j* in replicate *h*.

To estimate the contribution of variance in the per-deme means, we compute the variance across replicates for a given deme, and then take the average of these values across demes:

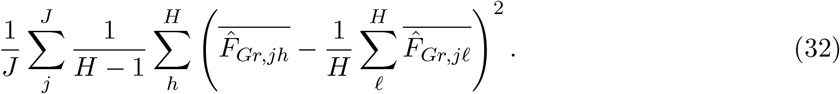

(here, the sums over 𝓁 and *h* are both sums over replicates–one for the mean, and one for the variance–but we use different letters to avoid confusion).

The denominator, in turn, can be estimated straightforwardly as

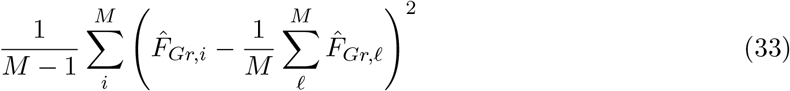

where we now use 𝓁 to index individuals within the mean calculation. Our estimate of the error is then given by summing (30) and (32) and dividing by (33).

#### 5.6.2 Principal components

To estimate the error in the sample PCs, we follow similar steps, except that it is not obvious how to compute the variance of the per deme means, as the relationship between the order of the underlying population PCs and the sample PCs may differ across replicates due to the noisiness of the sample PCs. We therefore include only the variance among individuals within demes in our estimate of the error, which makes it an estimate of a lower bound on the error, rather than a direct estimate. The PCs are automatically standardized to have a variance of 1, so that for the *k*^*th*^ PC, a lower bound on the error is given by

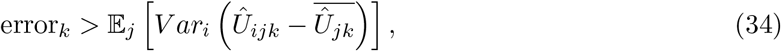

which we estimate as

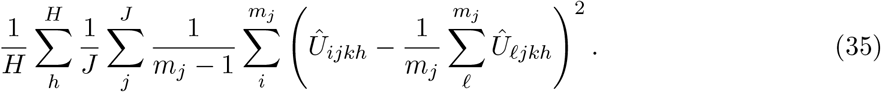

## Data availability

All of the code developed to produce the figures and simulations in this paper is available in the github repository: https://github.com/jgblanc/PGS-differences-confounding. We used the existing software plink2 https://www.cog-genomics.org/plink/2.0/, msprime https://tskit.dev/msprime/docs/stable/intro.html, bcftools https://samtools.github.io/bcftools/bcftools.html, R https://www.r-project.org/, and python https://www.python.org/.

## Acknowledgments

We would like to thank members of the Berg, Novembre and Steinrücken labs, as well as members of the University of Chicago genetics community for helpful discussions and feedback during the development of this project. We thank Andy Dahl and Matthew Stephens in particular for many helpful conversations. Additionally, we thank Matthew Stephens, John Novembre, and Xuanyao Liu for support at all stages of this work, as well as Maggie Steiner, Vivaswat Shastry, and Maryn Carlson for help troubleshooting and additional insights. Finally, we thank Graham Coop, Jeff Spence, Arjun Biddanda and Yuval Simons for comments on the manuscript. We also acknowledge funding support from the National Human Genome Research Institute (F31HG011821 to JGB) and the National Institute of General Medical Sciences (R35GM151257 to JJB).

---

**Figure S1:**
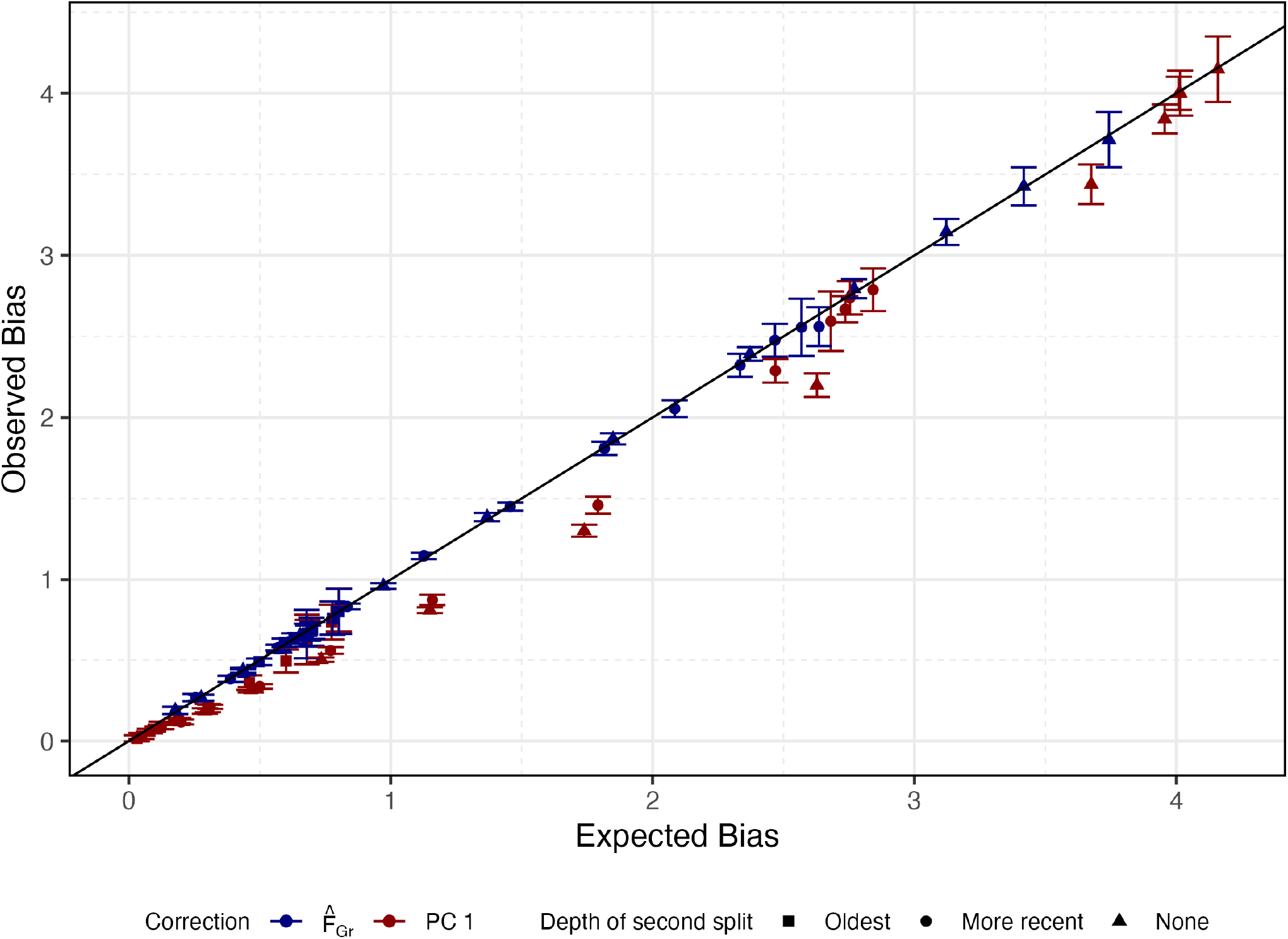
Error in estimates of 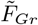 predicts bias in 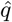 across population models. For all simulations in Figure 2 we compute the expected bias as 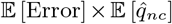 where 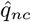 is the observed bias using effect sizes that were estimated with no correction. We then compare this expected bias to the observed bias when using that estimator as a covariate in the GWAS. The error in both 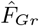 and sample PC 1 is highly predictive of the observed bias, though we observe that sample PC 1 exhibits a slight increase in bias reduction compared to the expected.

**Figure S2:**
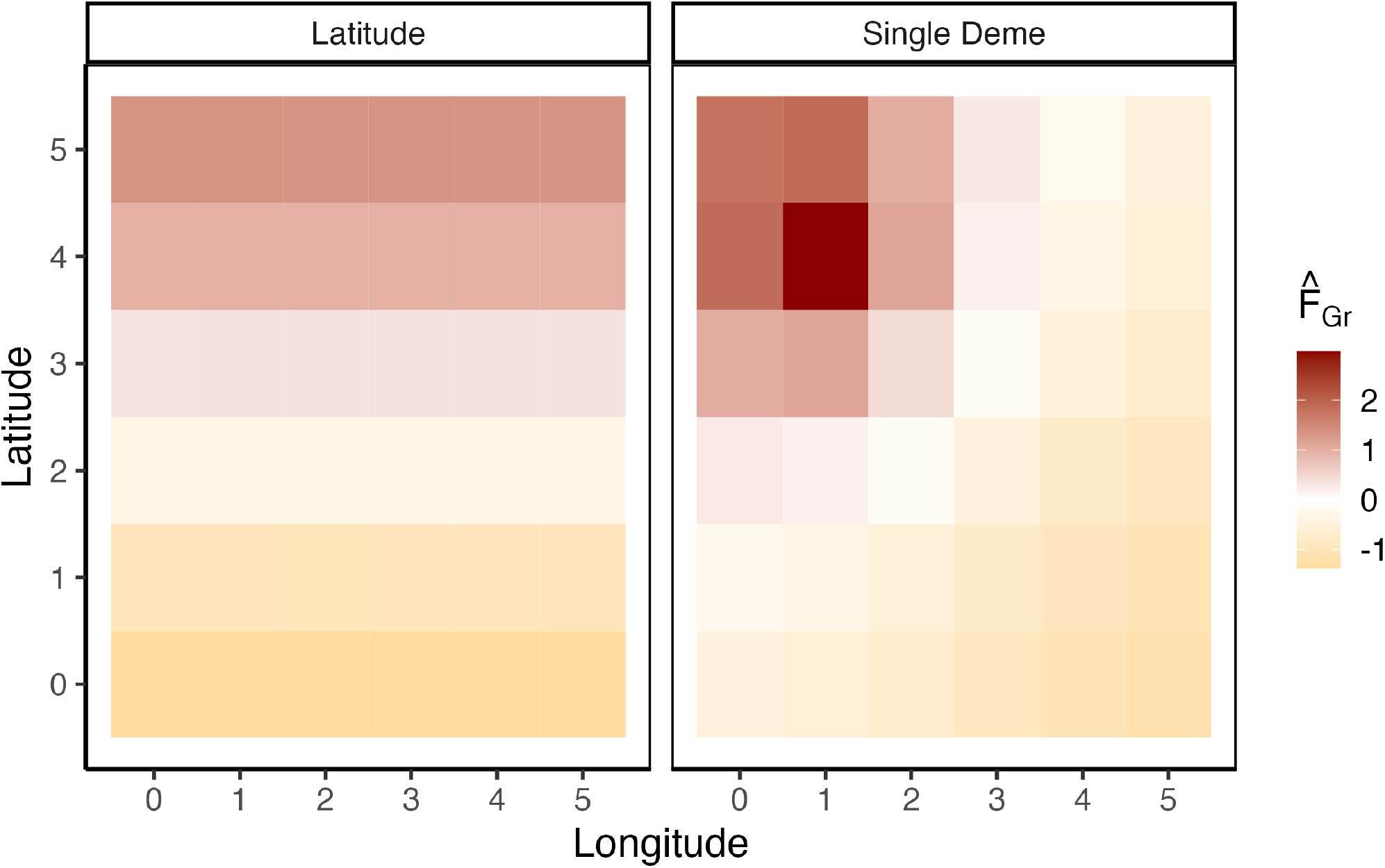
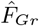 as observed in the GWAS panel. For both of the test vectors used in the grid simulations we plotted the average 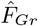 per deme across 100 replicates. For the latitudinal test vector, 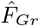 simply recapitulates latitude, which is unsurprising given the symmetric migration model we use. For the single deme test vector, 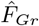 largely reflects the distance to the focal test deme.

**Figure S3:**
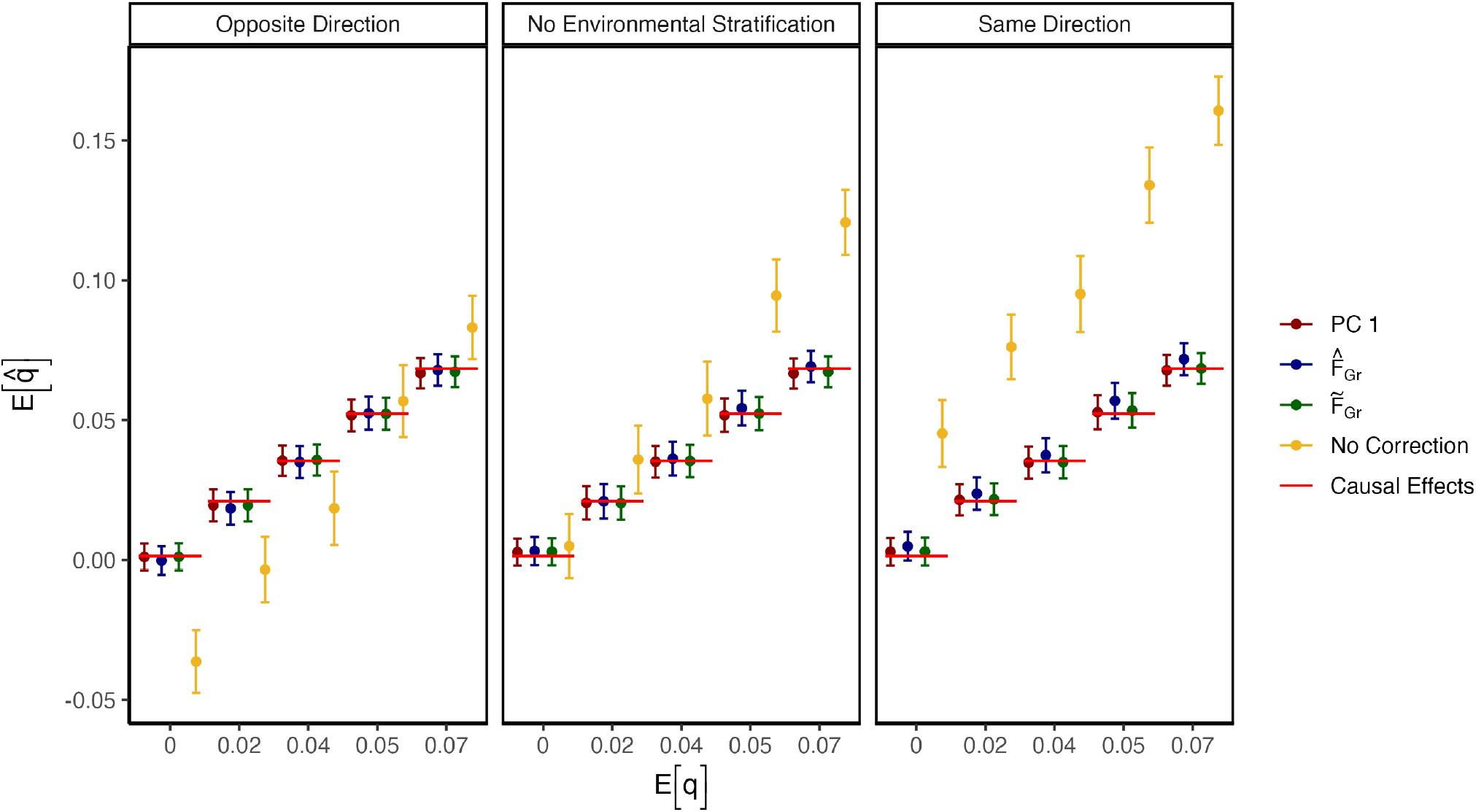
Including 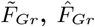, or PC 1 as a covariate in the GWAS model maintains power to detect true association signal. GWAS and test panels were simulated in the overlapping structure configuration (see Figure 1A). Heritable phenotypes (*h*^2^ = 0.3) were simulated with a true difference in polygenic scores by flipping the sign of a proportion of causal effects to align with allele frequency contrasts, *p*_*D,𝓁*_ − *p*_*C,𝓁*_, in the test panel. When stratification is in the same direction as the true difference, 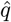 is upwardly biased, as it is when there is no environmental stratification, once genetic stratification is strong enough. When stratification is in the opposite direction, environmental and genetic stratification are opposed and the direction of bias depends on the strength of each. As expected, 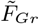 perfectly captures true association regardless of the direction of stratification. Estimators of 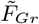 (i.e 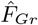 and PC 1) also capture true association, consistent with out theoretical arguments that downward bias is minimal when *S << L*.

