## supplementary text for "Testing for differences in polygenic scores in the presence of confounding"

June 25, 2024

**S2 Expectation of marginal effects** **6**

**S3 The expected polygenic scores** **7**

**S4 Polygenic score association test bias using marginal effects** **9**

**S5 Expectation of polygenic scores while controlling for $\tilde{\mathbf{F}}_{Gr}$** **10**

**S6 Polygenic score association test bias controlling for $\tilde{\mathbf{F}}_{Gr}$** **11**

**S7 Downward bias with true signal** **12**
S7.1 Expected bias . . . 12
S7.2 Toy model simulations . . . 13

**S8 Modeling the correlation structure among test panel individuals** **14**

### S1 Full Model

To model the distribution of genotypes in both panels, we assume that each individual's expected genotype at each site can be modelled as a linear combination of contributions from a potentially large number of ancestral populations, which are themselves related via an arbitrary demographic model. Natural selection, genetic drift, and random sampling each independently contribute to the distribution of genotypes across panels, and we make the approximation that these three effects can be combined linearly. We begin by first developing the population level model, before extending it to individuals.

#### S1.1 Population model

We assume that for each site $\ell$, the vector of allele frequencies across $K$ populations can be decomposed as

$$p_\ell = a_\ell + \Delta p_{\ell,D} + \Delta p_{\ell,S} \tag{S1}$$

where $a_\ell$ is the allele frequency in the original ancestral population, while the deviations $\Delta p_{\ell,D}$ and $\Delta p_{\ell,S}$ capture variation in allele frequency across populations due to genetic drift and natural selection respectively. We approximate the effect of drift via a multivariate Normal model, such that $\Delta p_{\ell,D} \sim MVN\left(0, a_\ell\left(1-a_\ell\right)\mathbf{F}_{pop}\right)$, where $\mathbf{F}_{pop}$ describes the covariance structure imposed by the population model [1, 2, 3, 4]. Selection induces an additional deviation of $\Delta p_{\ell,S} = \eta\beta_\ell a_\ell\left(1-a_\ell\right)A_{pop}$, where $\beta_\ell$ is site $\ell$'s effect (in standardized units) on the phenotype of interest, $\eta$ is the strength of selection on that phenotype, and $A_{pop}$ is a vector recording the extent to which each population inherits from the ancestral population in which selection occurred. Under the null, $\eta = 0$. Under the alternative, $\eta \sim N\left(0, \sigma_\eta^2\right)$, so that

$$\Delta p_{\ell,D} + \Delta p_{\ell,S} \sim MVN\left(0, a_\ell\left(1-a_\ell\right)\left(\mathbf{F}_{pop} + \sigma_\eta^2\beta_\ell^2 a_\ell\left(1-a_\ell\right)A_{pop}A_{pop}^\top\right)\right) \tag{S2}$$

where $A_{pop}A_{pop}^\top$ is a rank one matrix accounting for the impact of selection on the joint distribution of allele frequencies. We assume the trait is sufficiently polygenic and/or that selection is sufficiently weak that

$$\beta_\ell^2 a_\ell\left(1-a_\ell\right) \ll \frac{1}{\sigma_\eta^2} \quad \text{for all } \ell, \tag{S3}$$

so that for any individual variant, the effects of drift dominate over those of selection (i.e. $\Delta p_{\ell,D} >> \Delta p_{\ell,S}$ for all $\ell$). This condition is met, for example, if there are at least hundreds of loci and $\sigma_\eta^2$ is on the order of one.

#### S1.2 Sampling individuals for GWAS and test panels

Next, we consider taking two samples, one to compose the test panel and one to compose the GWAS panel. Individuals in each panel are created as mixtures of the underlying populations. There are $N$ test panel individuals and the deviation of their genotypes from the expected genotype in the ancestral population ($2a_\ell$) is

$$X_\ell = X_{\ell,D} + X_{\ell,S} + X_{\ell,B}, \tag{S4}$$

where $X_{\ell,D} = 2\mathbf{W}_X \Delta p_{\ell,D}$, $X_{\ell,S} = 2\mathbf{W}_X \Delta p_{\ell,S}$. The matrix $\mathbf{W}_X$ has dimensions $N \times K$, with rows specifying the fraction of ancestry that each of the $N$ test panel individuals inherit from each of the $K$ populations. We can think of the quantity $2a_\ell + X_{\ell,D} + X_{\ell,S}$ as giving a set of expected genotypes given the evolutionary history of the population, while $X_{\ell,B}$ contains the binomial sampling deviations across individuals given these expected genotypes.

Similarly, for the $M$ GWAS panels individuals, the deviation of their genotypes can be decomposed as

$$G_\ell = G_{\ell,D} + G_{\ell,S} + G_{\ell,B}, \tag{S5}$$

where $G_{\ell,D} = 2\mathbf{W}_G \Delta p_{\ell,D}$, $G_{\ell,S} = 2\mathbf{W}_G \Delta p_{\ell,S}$, $W_G$ is an $M \times K$ matrix with rows specifying the amount of ancestry that each GWAS panel individual inherits from each of the $K$ populations and $G_{\ell,B}$ captures the binomial sampling variance given the expected genotypes of the GWAS panel individuals (similar to above, $2a_\ell + G_{\ell,D} + G_{\ell,S}$ specifies this set of individual specific expected genotypes).

#### S1.3 Individual level model

Individuals in the two panels can draw ancestry from the same populations, or from related populations, which induces the joint covariance structure

$$Var\left(\begin{bmatrix} X_{\ell,D} \\ G_{\ell,D} \end{bmatrix}\right) = 4a_\ell\left(1 - a_\ell\right)\mathbf{F} \tag{S6}$$

where the matrix

$$\mathbf{F} = \begin{bmatrix} \mathbf{F}_{XX} & \mathbf{F}_{XG} \\ \mathbf{F}_{GX} & \mathbf{F}_{GG} \end{bmatrix} \tag{S7}$$

contains the within and between panel relatedness coefficients. These are in turn related to the population level covariance matrix as $\mathbf{F}_{XX} = W_X \mathbf{F}_{pop} W_X^\top$, $\mathbf{F}_{GG} = W_G \mathbf{F}_{pop} W_G^\top$, and $\mathbf{F}_{GX} = \mathbf{F}_{XG}^\top = W_G \mathbf{F}_{pop} W_X^\top$.

Similarly, the genotypic deviations due to selection can be written at the individual level as $X_{\ell,S} = 2\eta\beta_\ell a_\ell\left(1 - a_\ell\right) A_X$ and $G_{\ell,S} = 2\eta\beta_\ell a_\ell\left(1 - a_\ell\right) A_G$, where $A_X = \mathbf{W}_X A_{pop}$ and $A_G = \mathbf{W}_G A_{pop}$ describe the extent to which individuals in each of the panels inherit from the selection event.

The joint covariance structure of the individual genotypes is therefore

$$Var\left(\begin{bmatrix} X_\ell \\ G_\ell \end{bmatrix}\right) = 4a_\ell\left(1 - a_\ell\right)\left(\mathbf{F} + \sigma_\eta^2\beta_\ell^2 a_\ell\left(1 - a_\ell\right)\mathbf{A} + \mathbf{B}_\ell\right) \tag{S8}$$

where

$$\mathbf{A} = \begin{bmatrix} A_X A_X^\top & A_X A_G^\top \\ A_G A_X^\top & A_G A_G^\top \end{bmatrix} \tag{S9}$$

and $\mathbf{B}_\ell$ is a diagonal matrix with entries accounting for over-dispersion of the binomial sampling step due to evolutionary variation of the underlying allele frequencies (i.e. from drift and selection). The exact details of this matrix are not important other than that it is diagonal, i.e. it does not contribute to covaraince among individuals.

#### S1.4 Phenotypes

As in main text Equation (5), we assume that the vector of mean-centered phenotypes for the $M$ individuals in the GWAS panel can be written

$$\begin{aligned} y &= \sum_{\ell}^{S} \beta_\ell G_\ell + e \\ &= u + e \end{aligned} \tag{S10}$$

where $u = \sum_{\ell}^{S} \beta_\ell G_\ell$ is the combined genetic effect of all $S$ causal variants, and $e$ represents the combination of all environmental effects. The distribution of environmental effects is $e \sim MVN\left(c, \sigma_e^2 \mathbf{I}\right)$, where $c$ is the vector of expected environmental effects. We consider $c$ to be fixed (as opposed to random).

The genetic effect, $u$, can be broken down into the contributions from drift, selection, and sampling. In this case, $u_S = \sum_{\ell}^{S} \beta_\ell G_{\ell,S}$ is the vector of expected values of the genetic contributions to the phenotype, given the ancestries of the individuals in the GWAS panels. Both $u_D$ and $u_B$ have expectation zero and $u_D$ has a correlation structure determined by relatedness matrix $\mathbf{F}_{GG}$ (i.e $u_D \sim MVN(0, 2V_A \mathbf{F}_{GG})$), where $V_A = 2\sum_{\ell}^{S} \beta_\ell^2 a_\ell \left(1 - a_\ell\right)$ is the additive genetic variance in the original ancestral population. The full covariance structure of $u$ is therefore

$$Var(u) = 2V_A \mathbf{F} + \sigma_\eta^2 \left(V_A^2 + C_A^2\right) \mathbf{A} + 2V_A \mathbf{B}_u \tag{S11}$$

where $C_A = \sigma_\eta^2 \sum_{\ell \neq \ell'} \beta_\ell^2 \beta_{\ell'}^2 a_\ell \left(1 - a_\ell\right) a_{\ell'} \left(1 - a_{\ell'}\right) A_G A_G^\top$ and $\mathbf{B}_u = \sum_\ell \mathbf{B}_\ell$

We are ultimately interested in the behavior of polygenic scores given that some stratification effect exists in the phenotype. To this end, from here forward we will generally condition on the strength of selection on the phenotype, $\eta$, treating it as a fixed effect alongside $c$. Doing so gives us the condition expectations and covaraince structures:

$$\mathbb{E}\left[y \mid \eta, c\right] = u_S + c \tag{S12}$$

and

$$Var\left(y \mid \eta, c\right) = 2V_A \left(\mathbf{F} + \mathbf{B}_u\right) + \sigma_e^2 \mathbf{I}. \tag{S13}$$

#### S1.5 Polygenic scores

We consider a vector of mean centered polygenic scores, computed in the test panel. If the causal effects were known, then the contribution of the $S$ causal sites included in the polygenic score would be

$$Z = \sum_{\ell}^{S} \beta_\ell X_\ell. \tag{S14}$$

We continue now to condition on the effects of selection. Doing so, $Z$ is a vector of random variables with the randomness coming from $X_{\ell,D} + X_{\ell,B}$, neutral variation in the test panel genotypes due drift and binomial sampling, respectively. Therefore, the vector of expected polygenic scores for the $N$ individuals in the test panel is

$$\begin{aligned}\mathbb{E}\left[Z \mid \eta\right] &= \sum_{\ell}^{S} \beta_\ell \mathbb{E}\left[X_\ell \mid \eta\right] \\ &= \sum_{\ell}^{S} \beta_\ell X_{\ell,S} \\ &= \eta V_A A_X.\end{aligned} \tag{S15}$$

Here the expected polygenic scores depend on the strength of selection on the phenotype, the ancestral additive genetic variance, and the extent to which each individual in the test panel inherits from the selection event.

#### S1.6 Polygenic score association tests

We want to test the hypothesis that the polygenic scores are associated with some test vector, $T$, more than is expected due to drift alone. We do not necessarily assume that $T$ is equal to $A_X$, the axis along which selection has actually perturbed the polygenic scores.

To test for association of polygenic scores with the test vector, we consider the linear model

$$Z = qT + \varepsilon \tag{S16}$$

where $\varepsilon$ is i.i.d. Normal across individuals. A more powerful test is available by modeling the correlation structure among individuals, but the simpler i.i.d. model is sufficient for our purposes (see section S8). We assume that the test vector, $T$, is scaled to have a variance of one, so the slope is given by

$$q = \frac{1}{N} Z^\top T, \tag{S17}$$

and its conditional expectation by,

$$\begin{aligned}\mathbb{E}\left[q \mid \eta\right] &= \frac{1}{N} \mathbb{E}\left[Z \mid \eta\right]^\top T \\ &= \frac{1}{N} \eta V_A A_X^\top T\end{aligned} \tag{S18}$$

Under the null model, $\eta = 0$, so $\mathbb{E}[q] = 0$, reflecting the fact that genetic drift is directionless (though we may also have $\mathbb{E}\left[q\right] = 0$ if the test vector is not aligned with the axis along which selection has perturbed the polygenic scores, i.e. if $A_X^\top T = 0$). We compare this null hypothesis to an alternative in which $\eta \neq 0$ (assuming also that $A_X^\top T \neq 0$), reflecting the possibility that the polygenic score may have either a positive or negative association with the test vector. We refer to this as a "polygenic score association test".

We also re-frame this test as a statement about the association between the effect sizes and a set of genotype contrasts, $r_\ell = \frac{1}{N} X_\ell^\top T$, which measure the association between the test vector and the genotypes at each site [5]. Similar to the decomposition of the test panel genotypes above, we can decompose the genotype contrasts as

$$r_\ell = r_{\ell,D} + r_{\ell,S} + r_{\ell,B}, \tag{S19}$$

which capture contributions from drift, selection, and sampling, respectively. Writing $\beta$ and $r$ for the vectors of effect sizes and genotype contrasts across loci, we can write the test statistic as

$$q = \beta^\top r. \tag{S20}$$

#### S2 Expectation of marginal effects

Conditional on the GWAS panel genotypes and the genetic and environmental components of the phenotype, the marginal association at site $\ell$ is given by

$$\hat{\beta}_\ell \mid G_\ell, u, e = \frac{y^\top G_\ell}{G_\ell^\top G_\ell} \tag{S21}$$

$$= \beta_\ell + \frac{u_{-\ell}^\top G_\ell}{G_\ell^\top G_\ell} + \frac{e^\top G_\ell}{G_\ell^\top G_\ell} \tag{S22}$$

(equation (7) in the main text). Now, taking the expectation over the randomness due to drift and sampling in the GWAS panel genotypes and the total genetic effect, and over the random component of the environmental effect on the phenotype, the expected effect size estimate at site $\ell$ is

$$\mathbb{E}\left[\hat{\beta}_\ell \mid \eta, c\right] = \beta_\ell + \mathbb{E}\left[\frac{u_{-\ell}^\top G_\ell}{G_\ell^\top G_\ell} \mid \eta\right] + \mathbb{E}\left[\frac{e^\top G_\ell}{G_\ell^\top G_\ell} \mid c\right] \tag{S23}$$

$$= \beta_\ell + \mathbb{E}\left[u_{-\ell}^\top \mid \eta\right]\mathbb{E}\left[\frac{G_\ell}{G_\ell^\top G_\ell} \mid \eta\right] + \mathbb{E}\left[e^\top \mid c\right]\mathbb{E}\left[\frac{G_\ell}{G_\ell^\top G_\ell} \mid \eta\right] \tag{S24}$$

$$= \beta_\ell + u_{S,-\ell}^\top \mathbb{E}\left[\frac{G_\ell}{G_\ell^\top G_\ell} \mid \eta\right] + c^\top \mathbb{E}\left[\frac{G_\ell}{G_\ell^\top G_\ell} \mid \eta\right] \tag{S25}$$

$$\approx \beta_\ell + u_{S,-\ell}^\top \frac{\mathbb{E}\left[G_\ell \mid \eta\right]}{\mathbb{E}\left[G_\ell^\top G_\ell \mid \eta\right]} + c^\top \frac{\mathbb{E}\left[G_\ell \mid \eta\right]}{\mathbb{E}\left[G_\ell^\top G_\ell \mid \eta\right]} \tag{S26}$$

$$\approx \beta_\ell + u_{S,-\ell}^\top \frac{G_{\ell,S}}{4a_\ell(1-a_\ell)(1+\overline{F}_G)} + c^\top \frac{G_{\ell,S}}{4a_\ell(1-a_\ell)(1+\overline{F}_G)} \tag{S27}$$

$$\approx \beta_\ell + \eta A_G^\top V_{A,-\ell} \frac{\eta \beta_\ell A_G}{2(1+\overline{F}_G)} + c^\top \frac{\eta A_G}{2(1+\overline{F}_G)} \tag{S28}$$

$$\approx \beta_\ell + \frac{\eta^2 \beta_\ell V_{A,-\ell}}{2(1+\overline{F}_G)} + \frac{\eta c^\top A_G}{2(1+\overline{F}_G)} \tag{S29}$$

$$\approx \beta_\ell \left(1 + \frac{1}{2}\frac{\eta^2 V_{A,-\ell}}{1+\overline{F}_G}\right) + \frac{1}{2}\frac{\eta c^\top A_G}{1+\overline{F}_G} \tag{S30}$$

where $1 + \overline{F_G} = 1 + \frac{1}{M}\sum_{m=1}^{M} f_{mm}$ is the average level of self relatedness in the GWAS panel. The approximation at line (S26) comes from approximating the expectation of the ratio as the ratio of expectations. There is an additional approximation in line (S27) that comes from assuming that

$\mathbb{E}\left[G_\ell^\top G_\ell \mid \eta\right] = 4a_\ell(1-a_\ell)(1+\eta^2\beta_\ell^2 a_\ell(1-a_\ell)+\overline{F}_G) \approx 4a_\ell(1-a_\ell)(1+\overline{F}_G)$, consistent with our assumption that the effects of selection are small at the level of individual loci, relative to the effects of drift and sampling.

Therefore, consistent with results from [6], the expected marginal effect estimate for a given site is expected to be equal to the causal effect if the null holds (i.e. if $\eta = 0$). In contrast, under our alternative model, the focal site and the genetic background have a positive empirical covariance as they are both influenced by selection. As a result, the expected marginal effect is biased away from zero (i.e. further in the direction of the causal effect), and may be biased in either direction depending on whether the environmental confounder is positively or negatively associated with the axis along which selection has perturbed the focal allele. For the following analyses concerning bias in the distribution of polygenic scores and in polygenic score association tests (see section S3 and S4) we assume that the above biases in marginal effect estimates induced by selection are small relative to those induced by shared drift between the GWAS and test panels [7, 8, 9, 10, 5].

### S3 The expected polygenic scores

Given the marginal effect estimates, the vector of $N$ polygenic scores in the test panel is given by,

$$\hat{Z}^\top \mid \mathbf{G}, \mathbf{X}, u, e = \sum_{\ell=1}^{S} \hat{\beta}_\ell X_\ell^\top \tag{S31}$$

$$= \sum_{\ell=1}^{S} \beta_\ell X_\ell + \sum_{\ell=1}^{S} \frac{u_{-\ell}^\top G_\ell}{G_\ell^\top G_\ell} X_\ell^\top + \frac{e^\top G_\ell}{G_\ell^\top G_\ell} X_\ell^\top. \tag{S32}$$

We are interested in the expected polygenic scores in settings where both genetic and environmental confounders exist. We can write this expectation as

$$\mathbb{E}\left[\hat{Z}^\top \mid \eta, c\right] = \sum_{\ell=1}^{S} \mathbb{E}\left[\beta_\ell X_\ell \mid \eta\right] + \sum_{\ell=1}^{S} \mathbb{E}\left[\frac{u_{-\ell}^\top G_\ell}{G_\ell^\top G_\ell} X_\ell^\top \mid \eta\right] + \sum_{\ell=1}^{S} \mathbb{E}\left[\frac{e^\top G_\ell}{G_\ell^\top G_\ell} X_\ell^\top \mid \eta, c\right]. \tag{S33}$$

The first sum is the expected "true" polygenic scores given the non-neutral effects on the test panel genotypes ($\sum_{\ell=1}^{S} \mathbb{E}\left[\beta_\ell X_\ell \mid \eta\right] = \sum_{\ell=1}^{S} \beta_\ell X_{\ell,S} = \mathbb{E}[Z]^\top = \eta V_A A_X$) while the second sum captures contributions to the polygenic scores due to associations between the genotypes at individual focal sites and the genetic component of the phenotype from all other sites ($u_{-\ell} = \sum_{\ell' \neq \ell} \beta_{\ell'} G_{\ell'}$). The third captures contributions due to associations between the genotypes and the environmental component. We consider each of the second two in turn, beginning with the environment, which is more straightforward.

For each site $\ell$, we can write

$$\begin{aligned}
\mathbb{E}\left[\frac{e^\top G_\ell}{G_\ell^\top G_\ell} X_\ell^\top \mid \eta, c\right] &\approx \mathbb{E}\left[\frac{e^\top G_{\ell,D}}{(G_{\ell,D}+G_{\ell,B})^\top (G_{\ell,D}+G_{\ell,B})} X_{\ell,D}^\top \mid c\right] && \text{(S34)}\\
&\approx \mathbb{E}\left[e^\top \mid c\right] \mathbb{E}\left[\frac{G_{\ell,D} X_{\ell,D}^\top}{(G_{\ell,D}+G_{\ell,B})^\top (G_{\ell,D}+G_{\ell,B})}\right] && \text{(S35)}\\
&\approx \frac{c^\top}{M} \mathbb{E}\left[\frac{G_\ell X_\ell^\top}{(G_{\ell,D}+G_{\ell,B})^\top (G_{\ell,D}+G_{\ell,B})/M}\right] && \text{(S36)}\\
&\approx \frac{c^\top}{M} \mathbb{E}\left[\frac{G_{\ell,D} X_{\ell,D}^\top}{(G_{\ell,D}+G_{\ell,B})^\top (G_{\ell,D}+G_{\ell,B})/M}\right] = \frac{1}{M} c^\top \tilde{\mathbf{F}}_{GX} && \text{(S37)}\\
&\approx \frac{c^\top}{M} \frac{\mathbb{E}\left[G_{\ell,D} X_{\ell,D}^\top\right]}{\mathbb{E}\left[(G_{\ell,D}+G_{\ell,B})^\top (G_{\ell,D}+G_{\ell,B})/M\right]} = \frac{1}{M} \frac{c^\top \mathbf{F}_{GX}}{\left(1+\overline{F_G}\right)}. && \text{(S38)}
\end{aligned}$$

The approximation in line (S34) arises from assuming that $G_{\ell,S}$ and $X_{\ell,S}$ are small compared to $G_{\ell,D}$ and $X_{\ell,D}$, and can therefore be ignored (see S2). Alternately, (S34) is exact under the null that $\eta = 0$. In line (S38), we make a ratio of expectations approximation. $\tilde{\mathbf{F}}_{GX}$ is the expected kinship matrix computed on standardized genotypes, which is approximately equal to $\frac{\mathbf{F}_{GX}}{(1+\overline{F_G})}$, with the approximation being better if $\overline{F_G}$ is small. Summing across sites, we have

$$\sum_{\ell=1}^{S} \mathbb{E}\left[\frac{e^\top G_\ell}{G_\ell^\top G_\ell} X_\ell^\top \mid \eta, c\right] \approx \frac{S}{M} c^\top \tilde{\mathbf{F}}_{GX}. \tag{S39}$$

where $S$ is the number of causal sites.

For the contributions to the polygenic score arising from associations between genotypes and the genetic component of the phenotype, we have

$$\begin{aligned}
\mathbb{E}\left[\frac{u_{-\ell}^\top G_\ell}{G_\ell^\top G_\ell} X_\ell^\top \mid \eta\right] &\approx \mathbb{E}\left[\frac{u_{-\ell}^\top G_{\ell,D}}{(G_{\ell,D}+G_{\ell,B})^\top (G_{\ell,D}+G_{\ell,B})} X_{\ell,D}^\top \mid \eta\right] && \text{(S40)}\\
&\approx \mathbb{E}\left[u_{-\ell}^\top \mid \eta\right] \mathbb{E}\left[\frac{G_{\ell,D} X_{\ell,D}^\top}{(G_{\ell,D}+G_{\ell,B})^\top (G_{\ell,D}+G_{\ell,B})}\right] && \text{(S41)}\\
&\approx \frac{u_{S,-\ell}^\top}{M} \mathbb{E}\left[\frac{G_{\ell,D} X_{\ell,D}^\top}{(G_{\ell,D}+G_{\ell,B})^\top (G_{\ell,D}+G_{\ell,B})/M}\right] && \text{(S42)}\\
&\approx \frac{u_{S,-\ell}^\top}{M} \mathbb{E}\left[\frac{G_{\ell,D} X_{\ell,D}^\top}{(G_{\ell,D}+G_{\ell,B})^\top (G_{\ell,D}+G_{\ell,B})/M}\right] = \frac{1}{M} u_{S,-\ell}^\top \tilde{\mathbf{F}}_{GX} && \text{(S43)}\\
&\approx \frac{u_{S,-\ell}^\top}{M} \frac{\mathbb{E}\left[G_{\ell,D} X_{\ell,D}^\top\right]}{\mathbb{E}\left[(G_{\ell,D}+G_{\ell,B})^\top (G_{\ell,D}+G_{\ell,B})/M\right]} = \frac{1}{M} \frac{u_{S,-\ell}^\top \mathbf{F}_{GX}}{\left(1+\overline{F_G}\right)}, && \text{(S44)}
\end{aligned}$$

where $u_{S,-\ell} = \sum_{\ell' \neq \ell} \beta_{\ell'} G_{S,\ell'} = u_S - \beta_\ell G_{\ell,S}$. All sites in our model are unlinked, allowing us to treat $u_{-\ell}$ as independent from $G_\ell$.

Summing across sites, we have

$$\sum_{\ell=1}^{S} \mathbb{E}\left[\frac{u_{-\ell}^{\top} G_\ell}{G_\ell^{\top} G_\ell} X_\ell^{\top}\right] \approx \frac{1}{M} \sum_{\ell=1}^{S} u_{S,-\ell}^{\top} \tilde{\mathbf{F}}_{GX} \tag{S45}$$

$$\approx \frac{1}{M} \sum_{\ell=1}^{S} \left(u_S^{\top} - \beta_\ell G_{\ell,S}^{\top}\right) \tilde{\mathbf{F}}_{GX} \tag{S46}$$

$$\approx \frac{1}{M} \left(S u_S^{\top} - u_S^{\top}\right) \tilde{\mathbf{F}}_{GX} \tag{S47}$$

$$\approx \frac{S-1}{M} u_S^{\top} \tilde{\mathbf{F}}_{GX} \tag{S48}$$

$$\approx \frac{S}{M} u_S^{\top} \tilde{\mathbf{F}}_{GX}. \tag{S49}$$

Putting genetic and environmental contributions together, the expected polygenic scores are approximately

$$\mathbb{E}\left[\hat{Z}^{\top}\right] = \sum_{\ell=1}^{S} \mathbb{E}\left[\beta_\ell X_\ell\right] + \sum_{\ell=1}^{S} \mathbb{E}\left[\frac{u^{\top} G_\ell}{G_\ell^{\top} G_\ell} X_\ell^{\top}\right] + \sum_{\ell=1}^{S} \mathbb{E}\left[\frac{e^{\top} G_\ell}{G_\ell^{\top} G_\ell} X_\ell^{\top}\right] \tag{S50}$$

$$\approx \mathbb{E}\left[Z\right]^{\top} + \frac{S}{M}\left(u_S^{\top} + c^{\top}\right) \tilde{\mathbf{F}}_{GX}, \tag{S51}$$

where we have made the conditioning implicit in our notation for the sake of consistentcy with main text. We continue in this vein for the rest of this supplement: all expectations from here on should be interpretted as conditional on $\eta$ and $c$.

### S4 Polygenic score association test bias using marginal effects

The expected value of our test statistic, $\hat{q}$, follows straightforwardly from the expected values of the polygenic scores. We have

$$\mathbb{E}\left[\hat{q}\right] = \frac{1}{N} \mathbb{E}\left[\hat{Z}^{\top}\right] T \tag{S52}$$

$$\approx \mathbb{E}\left[q\right] + \frac{S}{NM}\left(\mu_S^{\top} + c^{\top}\right) \tilde{\mathbf{F}}_{GX} T = \mathbb{E}\left[q\right] + \frac{S}{NM}\left(\mu_S^{\top} + c^{\top}\right) \tilde{F}_{Gr}. \tag{S53}$$

The bias in $\hat{q}$ is therefore approximately equal to

$$\mathbb{E}\left[\hat{q} - q\right] \approx \mathbb{E}[\hat{q}] - \mathbb{E}\left[q\right] \tag{S54}$$

$$\approx \frac{S}{NM}\left(\mu_S^{\top} + c^{\top}\right) \tilde{\mathbf{F}}_{GX} T = \frac{S}{NM}\left(\mu_S^{\top} + c^{\top}\right) \tilde{F}_{Gr} \tag{S55}$$

where

$$\tilde{F}_{Gr} = \mathbb{E}\left[\frac{G_{\ell,D} r_{\ell,D}^{\top}}{(G_{\ell,D}+G_{\ell,B})^{\top}(G_{\ell,D}+G_{\ell,B})/M}\right] \tag{S56}$$

$$= \tilde{\mathbf{F}}_{GX} T \tag{S57}$$

$$\approx \frac{W_G \mathbf{F}_{pop} T_{pop}}{1 + \overline{F_G}}. \tag{S58}$$

Here S58 shows how the axis captured by $\tilde{F}_{Gr}$ in the GWAS panel is related to the underlying populations in our model, and to the pattern of population structure among them. The vector $T_{pop} = W_X^\top T$ has length $K$ and captures the axis of the test in terms of the underlying populations, while $\mathbf{F}_{pop}T_{pop}$ similarly has length $K$, and captures the extent to which genetic drift along the path to each population is associated with the axis of population structure identified by the test vector. $\mathbf{F}_{pop}T_{pop}$ therefore describes the axis of confounding in terms of populations. Multiplying this vector by $W_G$ then rotates this axis into the individual space of the GWAS panel to give $\tilde{F}_{Gr}$.

### S5 Expectation of polygenic scores while controlling for $\tilde{\mathbf{F}}_{Gr}$

Now, controlling for $\tilde{F}_{Gr}$, the marginal effects are

$$\hat{\beta}'_\ell \mid G_\ell, u, e = \beta_\ell + \frac{u_{-\ell}^\top \mathbf{P} G_\ell}{G_\ell^\top \mathbf{P} G_\ell} + \frac{e^\top \mathbf{P} G_\ell}{G_\ell^\top \mathbf{P} G_\ell} \tag{S59}$$

where $\mathbf{P} = \left(\mathbf{I} - \frac{1}{\|\tilde{F}_{Gr}\|}\tilde{F}_{Gr}\tilde{F}_{Gr}^\top\right)$

Thus, conditional on the variables in the GWAS panel, the polygenic scores are

$$Z^\top \mid \mathbf{G}, u, e = \sum_{\ell=1}^{S} \hat{\beta}'_\ell X_\ell^\top \tag{S60}$$

$$= \sum_{\ell=1}^{S} \beta_\ell X_\ell + \sum_{\ell=1}^{S} \frac{u^\top \mathbf{P} G_\ell}{G_\ell^\top \mathbf{P} G_\ell} X_\ell^\top + \sum_{\ell=1}^{S} \frac{e^\top \mathbf{P} G_\ell}{G_\ell^\top \mathbf{P} G_\ell} X_\ell^\top \tag{S61}$$

so the expected polygenic scores can be written as

$$\mathbb{E}\left[Z^\top\right] = \sum_{\ell=1}^{S} \mathbb{E}\left[\beta_\ell X_\ell\right] + \sum_{\ell=1}^{S} \mathbb{E}\left[\frac{u^\top \mathbf{P} G_\ell}{G_\ell^\top \mathbf{P} G_\ell} X_\ell^\top\right] + \sum_{\ell=1}^{S} \mathbb{E}\left[\frac{e^\top \mathbf{P} G_\ell}{G_\ell^\top \mathbf{P} G_\ell} X_\ell^\top\right]. \tag{S62}$$

Following the same steps as above (see S34, S40, and S45), we take the expectation of each of the above terms separately. The first term is again the expected "true" polygenic scores ($\sum_{\ell=1}^{S} \mathbb{E}\left[\beta_\ell X_\ell\right] = \sum_{\ell=1}^{S} \beta_\ell X_{\ell,S} = \mathbb{E}[Z]^\top$). Then we consider the association between the environment and the residual genotypes,

$$\mathbb{E}\left[\frac{e^\top \mathbf{P} G_\ell}{G_\ell^\top \mathbf{P} G_\ell} X_\ell^\top\right] \approx \frac{c^\top}{M} \mathbf{P} \mathbb{E}\left[\frac{G_{\ell,D} X_{\ell,D}^\top}{(G_{\ell,D}+G_{\ell,B})^\top \mathbf{P} (G_{\ell,D}+G_{\ell,B}) / M}\right] \tag{S63}$$

$$\approx \frac{c^\top}{M} \mathbf{P} \tilde{\mathbf{F}}_{GX} \tag{S64}$$

$$\approx \frac{c^\top}{M}\left(\tilde{\mathbf{F}}_{GX} - \frac{1}{\|\tilde{F}_{Gr}\|}\tilde{F}_{Gr}\tilde{F}_{Gr}^\top \tilde{\mathbf{F}}_{GX}\right) \tag{S65}$$

$$\approx \frac{c^\top}{M} \tilde{\mathbf{F}}_{GX}^{\perp \tilde{F}_{Gr}} \tag{S66}$$

where we make an additional approximation by ignoring the $\mathbf{P}$ in the denominator, which is reasonable so long as $\tilde{F}_{Gr}$ explains only a small fraction of the variance in the GWAS panel genotypes. Summing of the $S$ causal sites in the polygenic score,

$$\sum_{\ell=1}^{S} \mathbb{E}\left[\frac{e^\top \mathbf{P} G_\ell}{G_\ell^\top \mathbf{P} G_\ell} X_\ell\right] \approx \frac{S}{M} c^\top \tilde{\mathbf{F}}_{GX}^{\perp \tilde{F}_{Gr}} \tag{S67}$$

Turning to the contribution to the polygenic score arising from the association between focal site and the residual genetic background, the expectation is

$$\mathbb{E}\left[\frac{u_{-\ell}^\top \mathbf{P} G_\ell}{G_\ell^\top \mathbf{P} G_\ell} X_\ell^\top\right] \approx \frac{u_{S,-\ell}^\top}{M} \mathbf{P} \mathbb{E}\left[\frac{G_{\ell,D} X_{\ell,D}^\top}{(G_{\ell,D}+G_{\ell,B})^\top \mathbf{P} (G_{\ell,D}+G_{\ell,B})/M}\right] \tag{S68}$$

$$\approx \frac{u_{S,-\ell}^\top}{M}\left(\tilde{\mathbf{F}}_{GX} - \frac{1}{\|\tilde{F}_{Gr}\|}\tilde{F}_{Gr}\tilde{F}_{Gr}^\top \tilde{\mathbf{F}}_{GX}\right) \tag{S69}$$

$$\approx \frac{u_{S,-\ell}^\top}{M} \tilde{\mathbf{F}}_{GX}^{\perp \tilde{F}_{Gr}} \tag{S70}$$

where we again make the approximation of ignoring $\mathbf{P}$ in the denominator. Summing across the S sites,

$$\sum_{\ell=1}^{S} \mathbb{E}\left[\frac{u^\top \mathbf{P} G_\ell}{G_\ell^\top \mathbf{P} G_\ell} X_\ell\right] \approx \frac{1}{M}\sum_{\ell=1}^{S} u_{S,-\ell}^\top \tilde{\mathbf{F}}_{GX}^{\perp \tilde{F}_{Gr}} \tag{S71}$$

$$\approx \frac{S}{M} u_S^\top \tilde{\mathbf{F}}_{GX}^{\perp \tilde{F}_{Gr}} \tag{S72}$$

Putting the true polygenic scores and the residual genetic and environmental contributions together, the expected polygenic scores after controlling for $\tilde{F}_{Gr}$ are approximately,

$$\mathbb{E}\left[\hat{Z}\right] \approx \mathbb{E}[Z] + \frac{S}{M}\left(\mu_S^\top + c^\top\right) \tilde{\mathbf{F}}_{GX}^{\perp \tilde{F}_{Gr}}. \tag{S73}$$

### S6 Polygenic score association test bias controlling for $\tilde{\mathbf{F}}_{Gr}$

Following from the distribution of polygenic scores after controlling for $\tilde{\mathbf{F}}_{Gr}$, the expected association test statistic is,

$$\mathbb{E}[\hat{q}] \approx \mathbb{E}[q] + \frac{S}{M}\left(\mu_S^\top + c^\top\right) \tilde{\mathbf{F}}_{GX}^{\perp \tilde{F}_{Gr}} T \tag{S74}$$

$$\approx \mathbb{E}[q] + \frac{S}{M}\left(\mu_S^\top + c^\top\right)\left(\tilde{\mathbf{F}}_{GX} - \frac{1}{\|\tilde{F}_{Gr}\|}\tilde{F}_{Gr}\tilde{F}_{Gr}^\top \tilde{\mathbf{F}}_{GX}\right) T \tag{S75}$$

$$\approx \mathbb{E}[q] + \frac{S}{M}\left(\mu_S^\top + c^\top\right)\left(\tilde{F}_{Gr} - \frac{1}{\|\tilde{F}_{Gr}\|}\tilde{F}_{Gr}\tilde{F}_{Gr}^\top \tilde{F}_{Gr}\right) \tag{S76}$$

$$\approx \mathbb{E}[q] + \frac{S}{M}\left(\mu_S^\top + c^\top\right)\left(\tilde{F}_{Gr} - \tilde{F}_{Gr}\right) \tag{S77}$$

$$\approx \mathbb{E}[q] \tag{S78}$$

such that the expected bias is

$$\mathbb{E}\left[\hat{q} - q\right] \approx 0. \tag{S79}$$

Here we highlight a reasonable assumption we make regarding our ability to detect true signal when controlling for $\tilde{F}_{Gr}$. As discussed in the main text, including $\tilde{F}_{Gr}$ as a covariate removes any correlation between $\hat{\beta}$ and $r$ under the null hypothesis where $\eta = 0$. Notably, under the alternative hypothesis, our ability to detect signal is preserved as we use variation along other axes in the GWAS panel to estimate effect sizes which in turn can still be correlated with $r_S$ in the test panel. We therefore rely on the assumption that for site $\ell$ there is enough remaining variation in $G_\ell$ after regressing out $\tilde{F}_{Gr}$ to estimate $\hat{\beta}_\ell$. If all variation in $G_\ell$ lies along $\tilde{F}_{Gr}$, $y^\top \mathbf{P} G_\ell \approx 0$ and we will be unable to estimate $\hat{\beta}_\ell$. For example, if $\tilde{F}_{Gr}$ represented individuals on opposite sides of a population split this would only apply to sites with fixed differences. This is unlikely to be a concern in practice as $F_{ST}$ is low in human populations [11] and most variants will not be perfectly correlated with a single axis of structure. Therefore, it is reasonable to assume that it will still be possible to accurately estimate $\beta_\ell$, albeit with slightly larger standard error.

### S7 Downward bias with true signal

#### S7.1 Expected bias

The direct estimator approach (i.e computing $\hat{F}_{Gr}$ and including it as covariate in the GWAS) proposes to use the test panel genotype data twice: once when controlling for stratification in the GWAS panel, and a second time when testing for an association between the polygenic scores and the test vector. Shouldn't this remove the signal we are trying to detect? While the answer is yes, at least for naive applications, the effect will be small so long as the number of SNPs used to compute the correction is large relative to the number included in the polygenic score.

To see why, we can rewrite the regression eq. 23 from the main text in terms of a sum of the contribution from our focal site and all other sites as

$$y = G_\ell \beta_\ell + \left( \frac{L-1}{L} \hat{F}_{Gr,-\ell} + \frac{1}{L} \frac{G_\ell r_\ell}{G_\ell^\top G_\ell / M} \right) \omega + e \tag{S80}$$

where $\hat{F}_{Gr,-\ell}$ is the estimate of $\tilde{F}_{Gr}$ that one would obtain using all sites other than site $\ell$. Thus, because our $\hat{F}_{Gr}$ includes a contribution from the focal site, controlling for it induces a slight bias in the estimated effect size, the sign of which depends on the sign of $r_\ell$. If $r_\ell$ is positive, $\hat{\beta}_\ell$ has a slight negative bias, whereas if $r_\ell$ is negative, the bias will be positive. Similar effects are noted elsewhere in the statistical genetics literature, for example in correcting for PCs of gene expression data [12], and in the use of linear mixed models in GWAS [13], where it has been termed "proximal contamination". Assuming that the variance of $\frac{r_\ell}{\sqrt{G_\ell^\top G_\ell / M}}$ across sites included in the score is similar to those used to compute the correction, the product $r_\ell \hat{\beta}_\ell$ will be biased toward 0 by a factor of approximately $\left(1 - \frac{1}{L}\right)$ for each site, owing to the fact that the focal site contributes approximately $\frac{1}{L}$ of our estimate $\hat{F}_{Gr}$. Our test statistic is a sum over contributions of $r_\ell \hat{\beta}_\ell$ from $S$ independent sites, so including $\hat{F}_{Gr}$ as a covariate when estimating effect sizes induces a downward bias of approximately $\left(1-\frac{S}{L}\right)$, i.e.

$$\mathbb{E}\left[\hat{q} \mid q\right] \approx q\left(1-\frac{S}{L}\right). \tag{S81}$$

Notably, controlling for sample PCs of the GWAS panel genotype matrix will induce a similar effect if the sample PCs capture $\tilde{F}_{Gr}$. In either case, the downward bias should be small so long as sites used to compute the polygenic score are only a small subset of those used to estimate $\hat{F}_{Gr}$. While this picture is somewhat complicated by the existence of linkage disequilibrium in real populations, for human population samples imputed to common reference panels, the effective number of SNPs is typically on the order of at least half a million [14], suggesting that even for a polygenic score that included, for example, 10,000 SNPs, the downward bias should be no more than 2%. An important caveat is that some methods for computing polygenic scores allow for all or at least a substantial fraction of all SNPs genome wide to make non-zero contributions to the polygenic score [15, 16]. Further concern about downward biases in applications could likely be ameliorated via the "leave one chromosome out" scheme (we implement this approach in **??**, see **??** for details) commonly implemented in the context of linear mixed models [13, 17] or via iterative approaches that first aim to ascertain SNPs using a genome-wide estimate of $\hat{F}_{Gr}$ before re-estimating effects using an estimate of $\hat{F}_{Gr}$ computed from sites not in strong LD with any of the ascertained sites. Understanding the behavior of tests for polygenic score-ancestry associations in the context of these methods will require careful attention to the methods' assumptions.

#### S7.2 Toy model simulations

Next, we wanted to confirm that including $\hat{F}_{Gr}$ or $\hat{U}_1$ does not regress out true signals of polygenic score divergence, consistent with our theoretical argument above. To do this, we modified our simulations of the confounded topology from Figure 1 by adding causal loci to make the trait heritable, with $h^2 = 0.3$, and sampled the sign of the effect for these causal loci to generate a correlation between the effect and the frequency differences between populations C and D. This procedure generates a positive test statistic and is conceptually equivalent adding a selection event on the internal branch in the population phylogeny.

When there was no environmental stratification (Figure S3, middle panel), genetic stratification eventually created an upward bias in the uncorrected $\hat{q}$, as the strength of the divergence signal increased. Similarly, when we added environmental stratification in the same direction as the genetic stratification (Figure S3, right panel) the upward bias increased in magnitude. Finally, when we added environmental stratification in the opposite direction (Figure S3, left panel), environmental and genetic stratification oppose one another and the observed bias depended on the strength of each. We then observe that in all cases including $\tilde{F}_{Gr}$, $\hat{F}_{Gr}$, or $\hat{U}_1$ (here $\hat{F}_{Gr}$, and $\hat{U}_1$ are well estimated) as a covariate in the GWAS eliminates bias while still capturing the true association signal. This is expected in all situations for $\tilde{F}_{Gr}$ and when $S << L$ for $\hat{F}_{Gr}$ and $\hat{U}_1$.

#### S8 Modeling the correlation structure among test panel individuals

In the main text, we test for the association between polygenic scores and the test vector using the linear model,

$$Z = qT + \varepsilon \tag{S82}$$

where $\varepsilon$ is i.i.d Normal across individuals. If $T$ is scaled to have a variance of 1, the slope is given by,

$$q = \frac{1}{N} Z^\top T \tag{S83}$$

which we use as our test statistic of interest.

However, a more powerful test is available by modeling the correlation structure among individuals in the test panel. Under our genotypic model the expected pattern of covariance of genotypes at site among individuals in the test panel is,

$$Var\left(X_\ell\right) = \mathbb{E}[X_\ell X_\ell^\top] = 4a_\ell\left(1 - a_\ell\right)\left(\mathbf{F}^*_{XX}\right), \tag{S84}$$

where $\mathbf{F}^*_{XX} = \mathbf{F}_{XX} + \mathbf{B}_X$. $\mathbf{F}_{XX}$ contains the within panel relatedness coefficients and $\mathbf{B}_X$ is a diagonal matrix with entries specifying the amount of additional variance for each individual due to sampling.

Now, because the polygenic scores are sums across a large number of sites, under the null model the distribution of standardized polygenic scores is approximately multivariate Normal

$$\frac{Z}{\sqrt{2V_A}} \sim MVN\left(0, \mathbf{F}^*_{XX}\right) \tag{S85}$$

where $V_A = 2\sum_\ell^S \beta_\ell^2 a_\ell(1-a_\ell)$ is the additive genetic variance of the polygenic scores in the original ancestral population.

To test for evidence of an association, we compare this null model to an alternative in which polygenic scores are associated with $T$,

$$\frac{Z}{\sqrt{2V_A}} \sim MVN\left(T\gamma, \mathbf{F}^*_{XX}\right), \tag{S86}$$

where $\gamma$ is the slope. Here we assume that $T$ is mean centered and scaled such that $\mathbf{F}^{*-\frac{1}{2}}_{XX}T$ has a variance of one. With this scaling, the generalized least squares estimate of $\gamma$ is given by $\hat{\gamma} = {T^\top \mathbf{F}^{*-1}_{XX} Z}/{(N\sqrt{2V_A})}$, where the $\mathbf{F}^{*-1}_{XX}$ term accounts for the evolutionary non-independence of the individuals in the panel. Under the null hypothesis, $\hat{\gamma} \sim N\left(0, \frac{1}{N}\right)$.

To remove this dependence on the sample size, we let the test statistic be $q_x = \sqrt{N}\hat{\gamma}$, so that

$$q_x = \frac{Z^\top \mathbf{F}_{XX}^{*-1} T}{\sqrt{2NV_A}} \tag{S87}$$

$$= \frac{1}{\sqrt{2NV_A}} \sum_{\ell=1}^{S} \beta_\ell X_\ell^\top \mathbf{F}_{XX}^{*-1} T \tag{S88}$$

$$= \frac{1}{\sqrt{2NV_A}} \beta r_x. \tag{S89}$$

Under the null $q_x \sim N(0,1)$. This test statistic is related to Berg and Coop's $Q_X$ statistic [5, 18] in a straight-forward way. For a test of selection along a single axis of ancestry variation, $Q_X = q_x^2$. More generally, the $Q_X$ statistic combines multiple tests along different axes of ancestry variation, accounting for non-independence among axes and providing the appropriate multiple testing penalty.

As discussed above, $q_x$ is proportional to the generalized least squares estimate of $\gamma$ whereas $q$ in the main text is proportional to the ordinary least squares estimate. Therefore, $q_x$ accounts for covariance and heteroscedasticity in test panel genotypes caused by within test panel population structure. This results in a more powerful test, in precisely the same way that linear mixed models increase the power of association tests in GWAS by accounting for covariance in the phenotypes due to genetic relatedness.

To apply our correction procedure while accounting for the non-independence among test panel individuals, we can compute $r_{x,\ell} = X_\ell^\top \hat{\mathbf{F}}_{XX}^{*-1} T$ and plug $r_x$ into equation 22 in the main text instead of $r_\ell$. This version of $\hat{F}_{Gr}$ can again be included as a covariate in the GWAS and those corrected effect sizes, in conjunction with $r_x$ can then be used to compute $q_x$. Overall, we would expect this procedure to result in slightly stronger protection against stratification bias, as the $r_{x,\ell}$ will have a slightly lower variance than the $r_\ell$, and thus lead to a more accurate estimated $\hat{F}_{Gr}$, though we have not systematically investigated this.
